# Arginine methyltransferase PRMT1 equipoises trophoblast development to prevent early pregnancy loss

**DOI:** 10.1101/2025.09.30.679604

**Authors:** Purbasa Dasgupta, Rajnish Kumar, Soma Ray, Namrata Roy, Asef Jawad Niloy, Mounika Vallakati, Courtney Marsh, Sebastian J. Arnold, Soumen Paul

**Affiliations:** Department of Pathology & Laboratory Medicine, University of Kansas Medical Center, Kansas City, KS 66160, USA; Institute for Reproduction and Perinatal Research, University of Kansas Medical Center, Kansas City, KS 66160, USA and; Department of Obstetrics and Gynecology, University of Kansas Medical Center, Kansas City, KS 66160, USA; Institute of Experimental and Clinical Pharmacology and Toxicology, Faculty of Medicine University of Freiburg, Albertstrasse 25, 79104 Freiburg, Germany, and 4CIBSS-Centre for Integrative Biological Signalling Studies, University of Freiburg; Freiburg, D-79104, Germany

**Keywords:** PRMT1, Placenta, Trophoblast Stem Cells, Cytotrophoblast, Early Pregnancy Loss

## Abstract

1-2% of all human pregnancies suffer from idiopathic recurrent pregnancy loss (RPL) and underlying molecular causes are poorly understood. Here we show that defective Protein Arginine Methyltransferase 1 (PRMT1) function in trophoblast progenitors is a molecular cause for early pregnancy failure. PRMT1 is conserved in trophoblast progenitors and conditional deletion of PRMT1 in mouse trophoblast progenitors arrests placenta and embryonic development leading to lethality ∼E7.5. Remarkably, a subset of idiopathic RPL is associated with loss of PRMT1 in cytotrophoblast progenitors (CTBs). Experiments with human trophoblast stem cells (hTSCs), derived from these RPL-patients as well as PRMT1-depleted hTSCs revealed that PRMT1 is crucial for trophoblast progenitors self-renewal. Employing RNA-seq and CUT&RUN-sequencing in hTSCs, CTBs and primary mouse trophoblast progenitors we discover that PRMT1 promotes transcription of trophoblast stem-state regulators, like TEAD4 and MYBL2, by directly enriching histone H4 arginine 3 asymmetric di-methylation (H4R3Me2a) at their chromatin loci. PRMT1 is also essential for extravillous trophoblast (EVT) development during human placentation, while loss of PRMT1 in hTSCs spontaneously promotes syncytiotrophoblast (STB) differentiation. Our findings indicate that PRMT1 is an epigenetic governor that orchestrates mammalian trophoblast development and implicate the therapeutic potential of targeting the PRMT1-H4R3Me2a axis to mitigate early pregnancy loss.

## Introduction

RPL, defined as the loss of two or more consecutive pregnancies is a major health concern and 1-2% of all implantation-confirmed conceptions suffer from idiopathic (of unknown origin) RPL. Idiopathic RPLs are devastating to patients, and a better understanding of underlying mechanisms is necessary for better diagnosis, appropriate counseling and development of patient-specific treatment options. Inefficient placentation due to defective development of trophoblast cell lineages have been implicated as one of the major causes in idiopathic RPL^1^.

During first-trimester of pregnancy a human placenta develops extensive villous structure, in which self-renewing CTB progenitors establish the stem/progenitor cell compartment. Along with extensive self-renewal process, CTB progenitors of a first trimester placenta undergo two differentiation pathways, the villous and extravillous. The villous differentiation of CTBs takes place in floating villi, which floats freely into maternal blood. The mononuclear CTBs undergo cell cycle arrest and fuse to form multinucleated syncytiotrophoblasts, which establish the maternal-fetal exchange interface. In contrast, the extravillous differentiation of CTB takes place in anchoring villi, which anchor to the maternal uterus. In the anchoring villi, CTBs establish a column of proliferating CTB progenitors, known as column CTBs (cCTBs). cCTBs are the earliest cell types that arise during EVT development. Eventually cCTBs at the distal end of the trophoblast column differentiate into migratory invasive EVT cells, which invade into the maternal uterine compartment. A subset of EVTs remodels the uterine artery to increase blood flow at the uterine-placental interface for adequate nutrient supply to the developing fetus. These EVTs are known as endovascular EVTs^2–4^. The remaining invasive EVTs within the uterine interstitium comprise the interstitial EVTs^4^, which interact with uterine cells for adaptation of the maternal immune system and physiology to the developing placenta. During human pregnancy, the EVT invasion is precisely regulated both temporally and spatially. Spatially the invasion is confined to the endometrium, the first third of the myometrium, and associated spiral arteries. Temporally, EVT invasion is confined to early pregnancy. Thus, precise regulation of self-renewal of CTB progenitors and their coordinated differentiation to STBs and extravillous EVTs during first-trimester placental development is essential for successful progression of pregnancy^5–11^.

Defect in CTB self-renewal, as well as defective EVT development during early human pregnancy has been implicated in RPL. We showed that a subset of idiopathic RPL is associated with defective self-renewal of CTB progenitors due to impaired expression of TEA Domain Transcription Factor 4 (TEAD4)^12^. Recently, another study implicated defective expression of MYB Proto-Oncogene Like 2 (MYBL2) in RPL and showed that MYBL2 function is necessary for both CTB self-renewal and for EVT development^13^. Furthermore, global gene expression analyses identified that RPLs are associated with altered expression of many key genes that are essential for CTB self-renewal or EVT development during first-trimester of human pregnancy. However, molecular mechanisms that are involved in altered transcription of key regulators, such as TEAD4 and MYBL2, in RPL are elusive.

Protein arginine methyltransferase (PRMT) enzymes regulate gene transcription, RNA splicing and DNA damage repair and play critical roles in cell growth, differentiation and survival^14^. However, importance of protein arginine methylation and function of PRMT enzymes are seldom studied in the context of human placentation. The PRMT family consists of 9 members and is divided into three types depending on their product specificity. Type I includes PRMT1, 2, 3, 4, 6 and 8; type II includes PRMT5 and 9; and type III includes only PRMT7^15^. All PRMTs are able to catalyze the monomethylation of arginine, but type I and II catalyze the second methyl group asymmetrically (ADMA) and symmetrically (SDMA), respectively, on the guanidino group of the arginine^15^. Gene knockout studies in mouse showed that among all PRMT family members, only PRMT1 and PRMT5 are essential for embryonic development^16–18^. Mice with null alleles of PRMT2, 3, 6, 7, 8 and 9 are viable^19–24^, whereas PRMT4-null mice die at birth^25^. Although both PRMT1 and PRMT5 are essential for mouse embryonic development, the knockout embryos show different phenotypes. Loss of *Prmt5* function is early embryonic-lethal due to the abrogation of pluripotent cells in blastocysts. In contrast, *Prmt1*-global knockout (*Prmt1*-KO) mouse embryos die at early post-implantation stage, ∼embryonic day (E)7.0.

PRMT1 plays critical roles in cell growth, differentiation and survival by catalyzing ADMA^15^ on the arginine residues of its target proteins. PRMT1-mediated ADMA on arginine 3 residues on Histone H4 (H4R3Me2a) has been shown to influence gene transcription in multiple contexts^26-28^. H4R3Me2a modification on a chromatin locus promotes recruitment of other factors, such as Tudor domain-containing proteins TDRD3^29^ and SWI/SNF complex member SMARCA4^28,30^. H4R3Me2a enrichment also facilitates recruitment of CBP/p300 leading to the acetylation of various histone molecules^28,31^, instigating RNA Polymerase II (RNA POL II) recruitment leading to transcriptional activation. Thus, PRMT1-mediated H4R3Me2a modification acts as a molecular switch to promote transcription at PRMT1 target genes.

PRMT1 is conserved in mammals, however, the importance of PRMT1 or importance of protein arginine methylation in mouse and human placentation have never been studied. In this study, we show that PRMT1 expression is conserved in trophoblast progenitors and invasive trophoblasts of rodent and human placentae. In contrast, PRMT1 expression is reduced during STB development. We also provide evidence that a subset of idiopathic recurrent pregnancy loss is associated with loss of PRMT1 expression. We show that PRMT1 is essential for maintaining self-renewal ability in human trophoblast progenitors and for EVT development. In contrast, PRMT1 prevents premature STB development. Thus, our findings indicate that PRMT1 equipoises human trophoblast development. Our mechanistic analyses in hTSCs as well as in mouse and human primary trophoblast progenitors indicate that in an early post-implantation mammalian embryo PRMT1 regulates transcription of trophoblast progenitor-specific genes by directly modulating H4R3Me2a modification at their chromatin loci. Our findings suggest that a conserved PRMT1-mediated gene regulatory mechanism may be important to maintain self-renewal ability of trophoblast progenitors and progression of pregnancy in mammals, including humans.

## Results

### PRMT1 is abundantly expressed in trophoblast progenitors and invasive trophoblasts of developing rodent and human placentae

We hypothesized that PRMT1 is a conserved regulator for trophoblast lineage development during mammalian placentation. To examine this hypothesis, first we profiled *Prmt1* mRNA expression in trophoblast cells of developing mouse placenta. We performed single-cell RNA-sequencing (scRNA-seq) with mouse placentae from different developmental stages (E)7.0 to (E)12.5 and clustered different trophoblast cell types based on expression of marker genes (Fig. S1A, B). The scRNA-seq analyses indicated that in a developing mouse placenta *Prmt1* is most abundantly expressed in the cell cluster, which corresponds to trophoblast stem and progenitor cells (TSPCs) within the extraembryonic ectoderm/ectoplacental cone (ExE/EPC) region at E7.0 placenta primordium (Fig 1A). We also noticed *Prmt1* expression in junctional zone cells (Fig. 1A, Fig. S1A, B). Subsequent clustering of the E7.0 trophoblast progenitor cells identified two distinct populations, one expressing high levels of *Prmt1* (*Prmt1*_HIGH_) and the other with low abundance of *Prmt1* expression (*Prmt1*_LOW_) (Fig. 1B). The *Prmt1*_HIGH_ cells also highly express murine trophoblast stem cell markers *Eomes, Elf5, Cdx2, Id2 and Esrrb,* whereas, the *Prmt1*_LOW_ progenitors showed higher expressions of differentiated trophoblast markers, such as *Dlx3, Nr6a1, Prl3b1, Tpbpa* and *Plac1* (Fig. 1C). Thus, during mouse placentation *Prmt1* mRNA is most abundantly expressed in TSPCs, which maintain the trophoblast stem-state gene expression program. Immunofluorescence studies in histological sections of E7.0 mouse and E9.0 rat conceptuses further confirmed abundant PRMT1 protein expression in TSPCs within both mouse and rat placenta primordia (Fig 1D,E). Interestingly, PRMT1 expression is strongly repressed in the developing labyrinth zone containing the STBs of an E9.5 mouse placenta (Fig. S2A). In contrast, we detected PRMT1 protein expression in the junctional zone of an E9.5 mouse placenta (Fig S2A), which contains the invasive trophoblast progenitors, and in the invasive trophoblast cells at E18.5 rat uterine-placental interfaces (Fig S2B).

**Figure 1.**
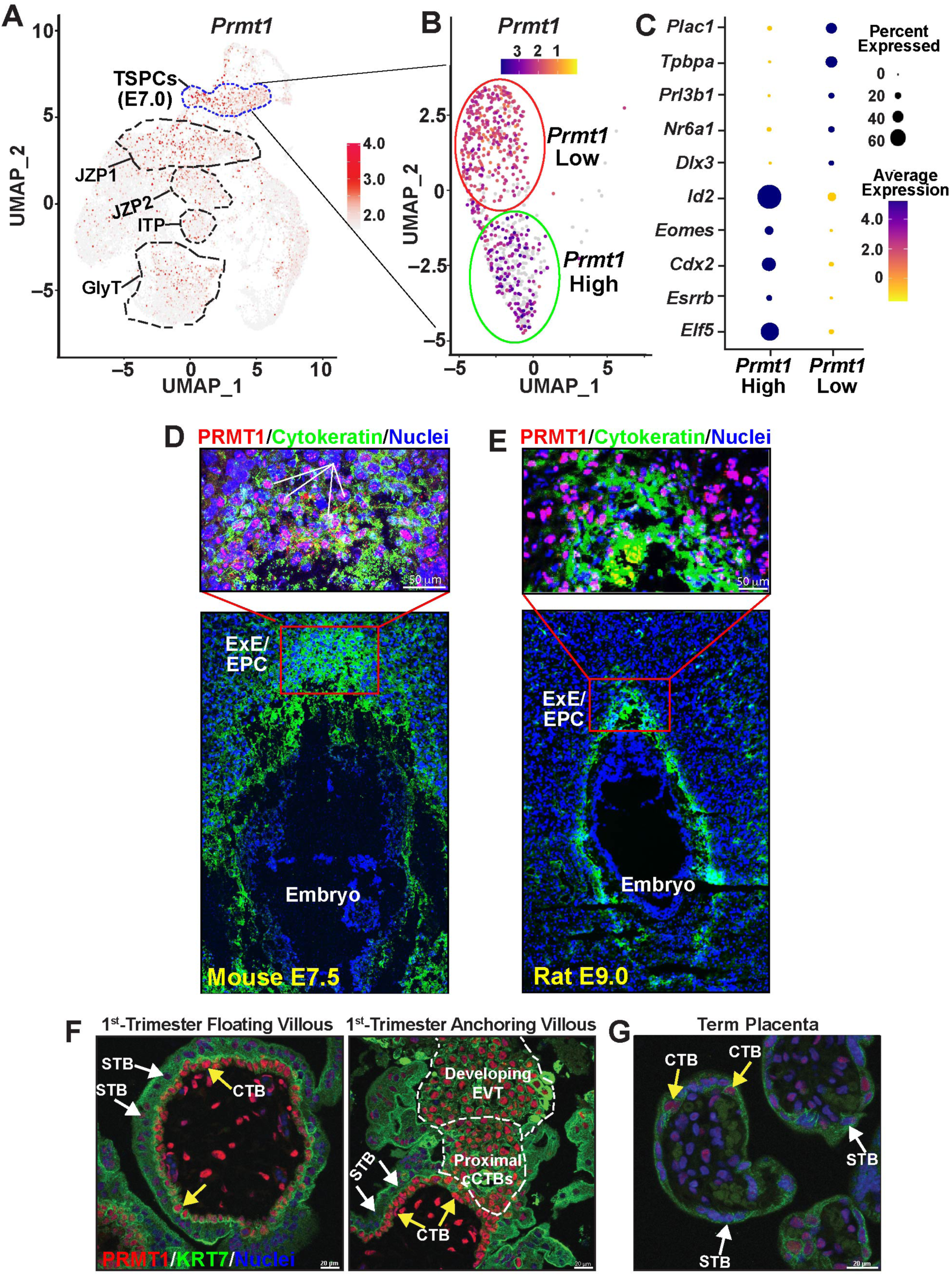
PRMT1 expression is conserved in rodent and human trophoblast progenitors. **(A)** scRNA-seq clusters of developing mouse placenta from E7.0 to E12.5 showing different cell clusters with *Prmt1* expressions. Note high *Prmt1* expression in cell cluster representing TSPCs from E7.0 developing placenta primordium. **(B)** Further clustering showing *Prmt1*_HIGH_ and *Prmt1*_Low_ cells within the TSPC population. **(C)** Dot plot showing average expression and percent of cells in *Prmt1*_HIGH_ and *Prmt1*_low_ TSPC clusters for genes representing undifferentiated TSC and differentiated trophoblast cell markers. **(D)** And **(E)** ExE/ EPC regions in an E7.0 mouse placenta primordium and E9.0 rat placenta primordium, respectively, showing PRMT1 (red) and Cytokeratin 7 (PanCK, green) expressions in TSPCs. **(F)** Histological section of a human first-trimester (week 7) placenta showing high level of PRMT1 expression (red) in CTBs of a floating villous (left) and in both proximal cCTB and distal EVT precursors within an anchoring villous (middle). Cytokeratin 7 (KRT7) was used to mark trophoblast cells. **(G)** Histological section of a term human placenta showing PRMT1 expression in trophoblast cells (right).

Next, we tested PRMT1 expression in trophoblast cells of a first-trimester human placenta. In a developing first trimester human placenta the proliferating villous CTBs comprise the stem/progenitor compartment and resembles with mouse TSPCs as they both express key conserved transcription factors, like CDX2, ELF5, TEAD4, GATA2 and GATA3^12,32,33^. We found that PRMT1 is highly expressed in proliferating CTBs within a floating villous, whereas expression is repressed in differentiated STBs (Fig.1F, Left)). In the anchoring villi, PRMT1 is highly expressed in column CTBs as well as in nascent EVTs, which develops from the distal end of the CTB column (Fig 1F, Right). We also confirmed similar expression patterns of *PRMT1* mRNA by reanalyzing recently published scRNA-seq data from first-trimester human placentae^34,35^ (Fig. S2C,D). We noticed that *PRMT1* mRNA is highly expressed in undifferentiated CTB and mature EVT cell clusters but is repressed within the committed STB cell cluster. In contrast to a 1^st^ trimester human placenta, PRMT1 expression significantly diminishes in the CTBs of a term placenta (Fig 1G), indicating that PRMT1 might have an essential role during early stages of human placentation.

Collectively, our studies indicated a conserved PRMT1 expression pattern during trophoblast development. PRMT1 is highly expressed in TSPCs of a developing rodent placenta and in CTB progenitors of a developing 1^st^ trimester human placenta. PRMT1 expression is maintained during invasive trophoblast/EVT development. However, STB development during both mouse and human placentation is associated with reduced PRMT1 expression.

### PRMT1 function in trophoblast progenitors of an early post-implantation mouse embryo is essential for placentation

An earlier report^16^ using a gene-trap allele showed that global loss of *Prmt1* in mouse embryo results in embryonic death ∼E6.5-E7.0, a stage when the extraembryonic placenta primordium, containing the ExE/EPC, develops. As PRMT1 is abundantly expressed in TSPCs, we tested whether TSPC development is affected in *Prmt1-KO* ExE/EPC regions. To that end, we studied *Prmt1^tm1a(EUCOMM)Wtsi^* mouse, that harbors the Knock-out first (Tm1a) allele (*Prmt1^tm1a^ mice),* developed by the Knockout Mouse Project^36^. Similar to the earlier report, we also noticed that homozygous *Prmt1^tm1a^ (PRMT1-KO)* embryos don’t develop beyond E7.0 (Figs 2A,B and S3A). Remarkably, the placenta primordium of ∼E7.0 *Prmt1-KO* conceptuses were smaller (Fig. 2B) and showed drastic loss of ESRRB-expressing TSPC population (Fig. 1C). Therefore, we next performed loss-of-function studies with *Prmt1*-conditional allele (*Prmt1^fl/fl^*) containing mice to test the importance of TSPC-specific PRMT1 function during early post-implantation mouse development.

**Figure 2:**
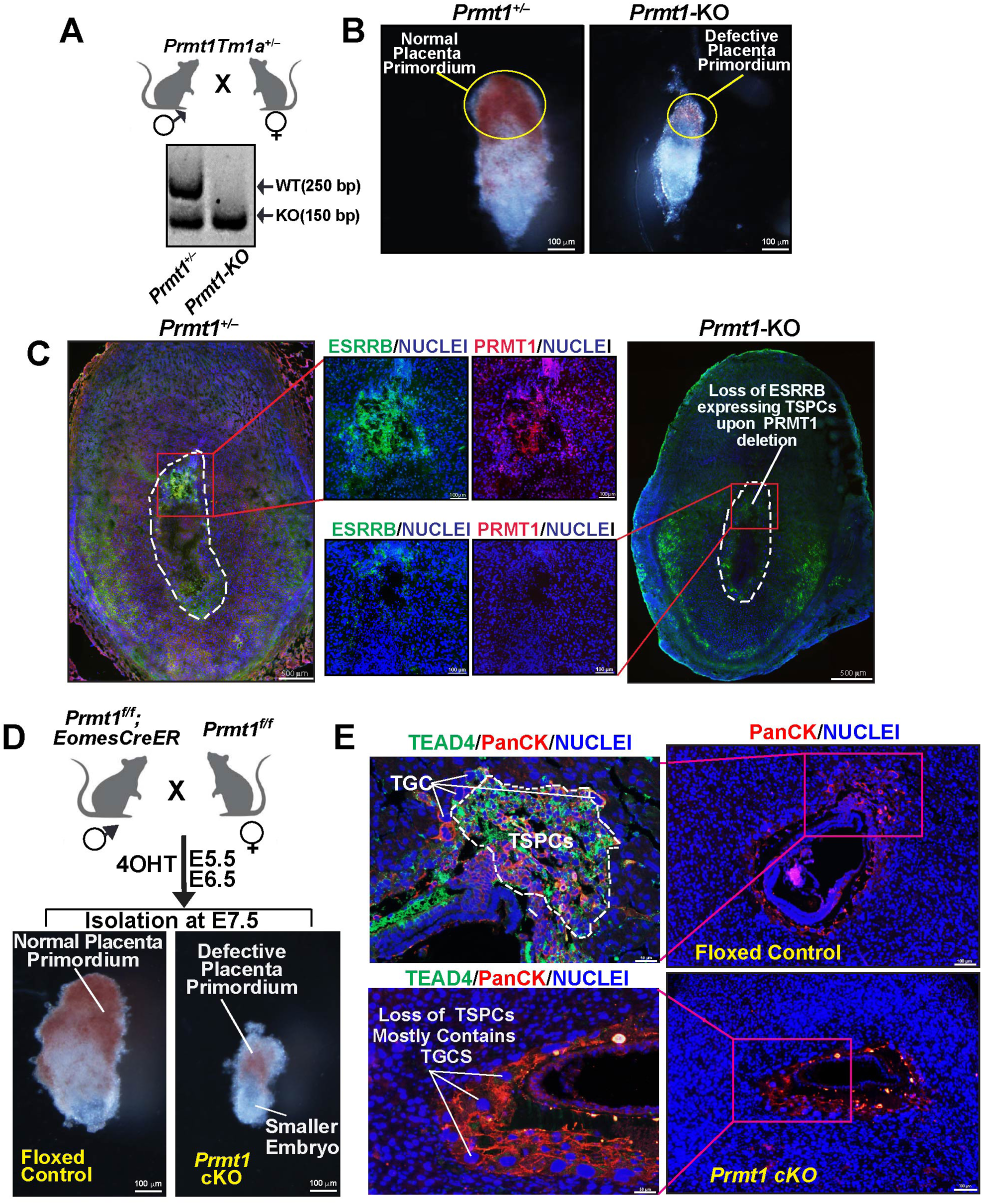
Loss-of PRMT1 in TSPCs of an early post-implantation mouse embryo impairs placentation leading to embryonic death. **(A)** Genotyping confirming *Prmt1* knockout allele in E7.0 *Prmt1*-*Tm1a* homozygous embryos. **(B)** Representative images of E7.0 *Prmt1-*heterozygous and *Prmt1-*KO conceptuses are shown. Embryonic and placentation defect were prominent in *Prmt1-* KO conceptus. **(C)** Immunostaining images of E7.0 *Prmt1-Tm1a* heterozygous (control, Left) and homozygous *Prmt1-KO* (right) showing ESRRB (green, TSPC marker) and PRMT1 (red) expression. The drastic loss of ESRRB expressing TSPCs at the ExE/EPC region of the *Prmt1-KO* conceptus is shown in the inset. **(D)** Conditional *Prmt1-alleles* were deleted in TSPCs of a developing mouse placenta and embryonic and placental developments were analyzed at E7.5 in Control *(Prmt1^fl/fl^)* and *Prmt1-cKO* embryos. Representative images of control and *Prmt1-c*KO conceptuses are shown. Embryonic and placentation defect were prominent in *Prmt1-c*KO conceptus. **(E)** TSPCs in the developing placenta primordium of control and *Prmt1-c*KO implantation sites were analyzed at ∼E7.5 via immunostaining with TEAD4 (green, TSPC marker) and pan-cytokeratin antibody (red, pan trophoblast marker). The developing *Prmt1-c*KO placenta primordium is smaller and mainly contains TGCs, which can be identified via larger nuclei (Blue).

We established a *Prmt1^fl/fl^:Eomes^CreER^* mouse strain by crossing *Prmt1^fl/fl^* mice with *Eomes^CreER^* mice, in which a tamoxifen-activatable Cre-recombinase (*CreER*) is expressed under the control of the Eomes genomic locus ^37^. Thus, *Prmt1* is conditionally deleted in TSPCs of *Prmt1^fl/fl^:Eomes^CreER^*conceptuses in presence of the synthetic estrogen receptor ligand, 4-hydroxytamoxifen (4-OHT). We crossed *Prmt1^fl/fl:^Eomes^CreER^*males with *Prmt1^fl/fl^* females to confine *CreER*-expression within the developing conceptus (Fig. 2D). We induced *Prmt1* gene deletion (Fig. S3B) starting at ∼E5.5 and monitored embryonic development in *Prmt1*-conditional knockout *(Prmt1-cKO)* conceptuses at *∼*E7.5. Conditional knockout of *Prmt1,* starting at ∼E5.5, resulted in smaller ExE/EPC region and also affected the embryonic development (Fig. 2D, Fig.S3C). Analyses of histological sections showed that placenta primordium of *Prmt1-cKO* conceptuses had drastically reduced TSPC population and contained mainly trophoblast giant cells (TGCs) (Fig. 2E). Furthermore, *ex vivo* cultures of the developing ExE/EPC regions revealed that deletion of PRMT1 impairs expansion of primary TSPCs (Fig. 2E, Fig. S3D), along with loss of transcription of TSPC-specific genes, such as *Tead4* (Fig. S3F). Collectively, our studies in *Prmt1*-cKO mouse established that PRMT1 is critical to promote self-renewal of TSPCs and progression of placenta/embryonic development in an early post-implantation mammalian embryo.

### Idiopathic RPL is associated with loss of PRMT1 expression in CTBs and developing EVTs

Idiopathic RPL is often associated with defective trophoblast development and placentation. Others and we have reported that idiopathic RPLs are associated with defective expression of key regulators of CTB self-renewal, such as TEAD4 and MYBL2^12,13^. In the mouse, TSPC-specific deletion of *Prmt1* prevents progression of embryonic development ∼E7.5, a developmental stage equivalent to the Carnegie stage 7 (days 15–17 post-conception) in human, when villous placenta rapidly develops with formation of primary and secondary villi due to extensive proliferation of CTB progenitors and their differentiation to STBs ^38^. These findings led us to hypothesize that defects in PRMT1 may contribute to early pregnancy loss in humans. Therefore, we collected placentae from patients with idiopathic RPLs (RPL-Placenta, Supplementary Table S1). We ensured that the collected placental tissues were not necrotic. We selected three first-trimester (<12.4 weeks) RPL placentae and one early 2^nd^ trimester (14+5 week) RPL-Placenta with visible defect in placental villi development (Fig. 3A) and performed global RNA-sequencing (RNA-seq) analyses (Fig. 3B, Supplementary Dataset S1). Similar to earlier reports^12,13,39^, RNA-seq analyses showed that mRNA expressions of *TEAD4* and *MYBL2* were downregulated in RPL-Placentae. In contrast, STB-specific genes, such as placenta specific glycoproteins (*PSGs*) and chorionic somatomammotropin hormones (*CSH1/2*), were induced in RPL-Placentae (Fig. 3B, Supplementary Dataset S1). Strikingly, we also observed strong loss of *PRMT1* mRNA expression in RPL-Placentae (Fig. 3B, Supplementary Dataset S1). Therefore, we tested PRMT1 protein expression in histological sections of RPL-placentae and confirmed reduced expression of PRMT1 in CTBs (Fig 3C, D). A subsets of RPL placentae showed significantly reduced PRMT1 expression in CTBs, whereas, some RPL placentae showed almost undetectable PRMT1 expression in CTBs. We also observed strong repression of PRMT1 expression in the CTB column of anchoring villi (Fig. 3E, F) in RPL-Placentae. Taken together, our studies confirmed that idiopathic RPL is often associated with loss of PRMT1 expression in CTB progenitors and in developing EVT precursors.

**Figure 3.**
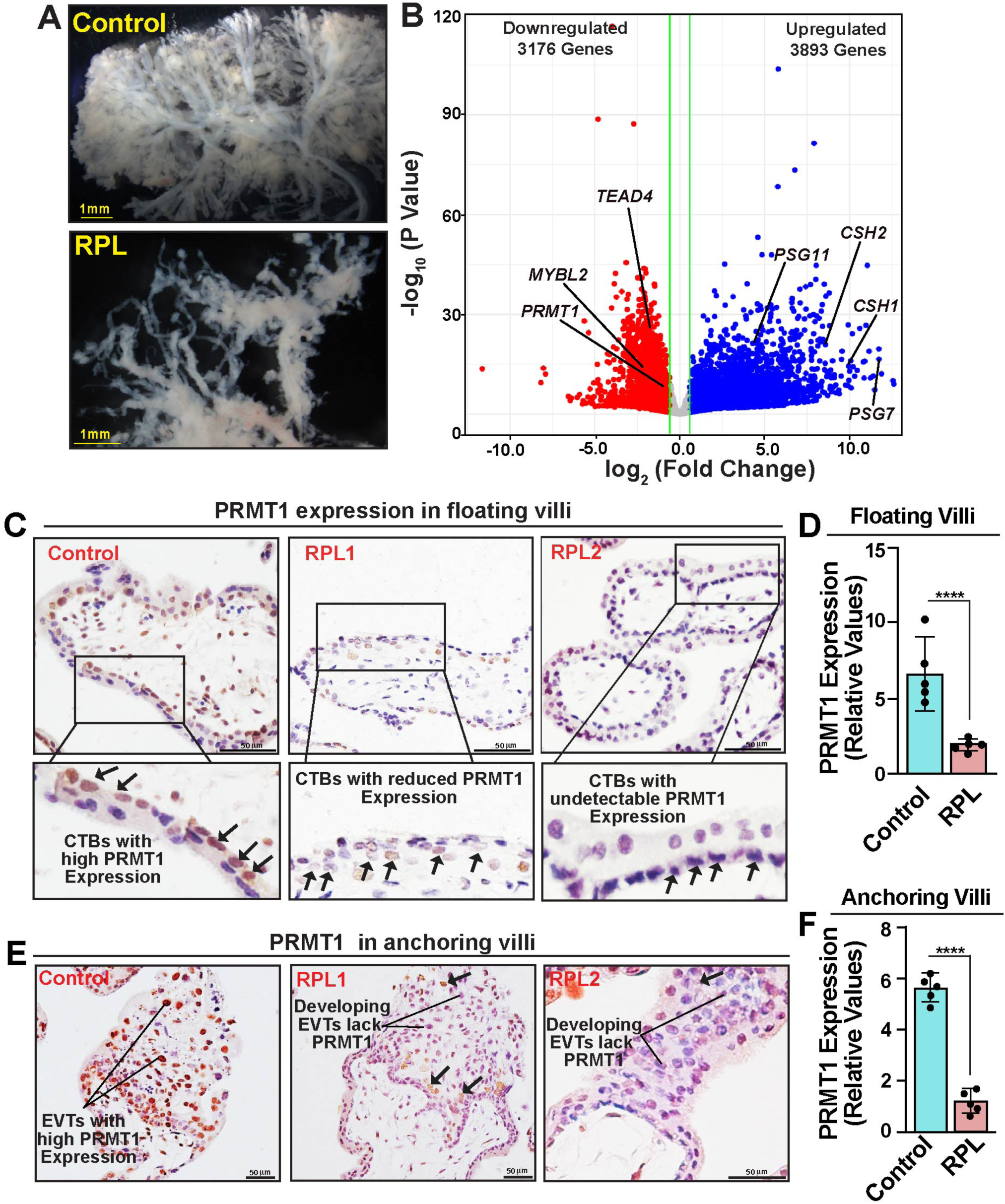
Idiopathic RPLs is associated with reduced PRMT1 expression in CTB progenitors and EVT Precursors. **(A)** Micrographs show defective villi formation in a first-trimester (9-week) placenta from a patient suffering from idiopathic RPL. An 8.5-week placenta from sporadic miscarriage without apparent defect in placental villi formation was used as a control. **(B)** Volcano plot showing global gene expression changes in RPL placentae with defect in placental villi formation. Unbiased RNA-seq analyses were performed and ≥1.5-fold gene-expression changes in RPL placentae with a false discovery rate of P < 0.05 were indicated with colored dots (blue: up-regulated, red: down-regulated). Significant down-regulation in expression of *TEAD4 (*a marker of undifferentiated CTBs*), MYBL2 (*important for CTB and EVT development*), PRMT1,* and up-regulation of *CSH1, CSH2, PSG7* and *PSG11* (STB-specific genes) are indicated. **(C)** Immunostaining images confirming strong reduction of PRMT1 protein expression in CTBs in floating villi of idiopathic RPL placentae. **(D)** Image J quantitation of PRMT1 protein expression in CTBs of control and Idiopathic RPL placentae (n=5 placentae for each condition, p<0.01). **(E)** Immunostaining images confirming reduced PRMT1 protein expression in proximal cCTBs and distal EVT precursors in anchoring villi of idiopathic RPL placentae. **(F)** Image J quantitation of PRMT1 protein expression in EVT precursors of control and idiopathic RPL placentae (n=5 placentae for each condition, p<0.01).

### PRMT1 is essential for maintaining self-renewal in human trophoblast progenitors

As idiopathic RPL is often associated with reduced PRMT1 expression in CTBs and EVT precursors, we next examined the consequences of silencing of PRMT1 in human trophoblast development. To that end we used hTSCs, which are derived from first-trimester CTBs^40^ as a model system. The established hTSCs can be maintained in a self-renewing stem state for multiple passages without affecting their self-renewal ability or without inducing genomic instability^12^. They can be efficiently differentiated to both STBs and EVTs. We confirmed that PRMT1 expression pattern in hTSCs robustly recapitulates *in vivo* expression patterns of PRMT1 (Fig. S4A). Immunofluorescence analyses revealed that PRMT1 is highly expressed in self-renewing hTSCs, and when they are differentiated to EVTs (Fig. S4A). In contrast, STB differentiation is associated with significant reduction in PRMT1 expression.

To perform loss-of-function analyses, we initially employed a CRISPR gene editing approach. However, we could not grow hTSCs after CRISPR-mediated gene deletion, indicating that PRMT1 is essential for the survival of hTSCs. We reasoned that there might not be any human pregnancy, in which PRMT1 is completely lost. Rather, pathological pregnancies are most likely associated with an alteration in the degree of PRMT1 expression in trophoblast cells. Therefore we employed an RNAi approach and specifically depleted PRMT1 in hTSCs by using lentiviral-mediated transduction of shRNAs (Fig. 4A, B). Expressions of other PRMT family were not significantly altered in PRMT1 Knock-down (*PRMT1-KD*) hTSCs (Fig. S4B). However, the loss of PRMT1 strongly reduced global ADMA modifications in hTSCs (Fig. 4C), confirming that PRMT1 is the major arginine methyl transferase for ADMA modification in human trophoblast cells. The *PRMT1-KD* hTSCs showed strong defect in cell proliferation (Fig. 4D, E) without significantly inducing cell death (Fig. S4C). The loss of cell proliferation was further confirmed via BrDU incorporation assay (Fig. 4F,G).

**Figure 4.**
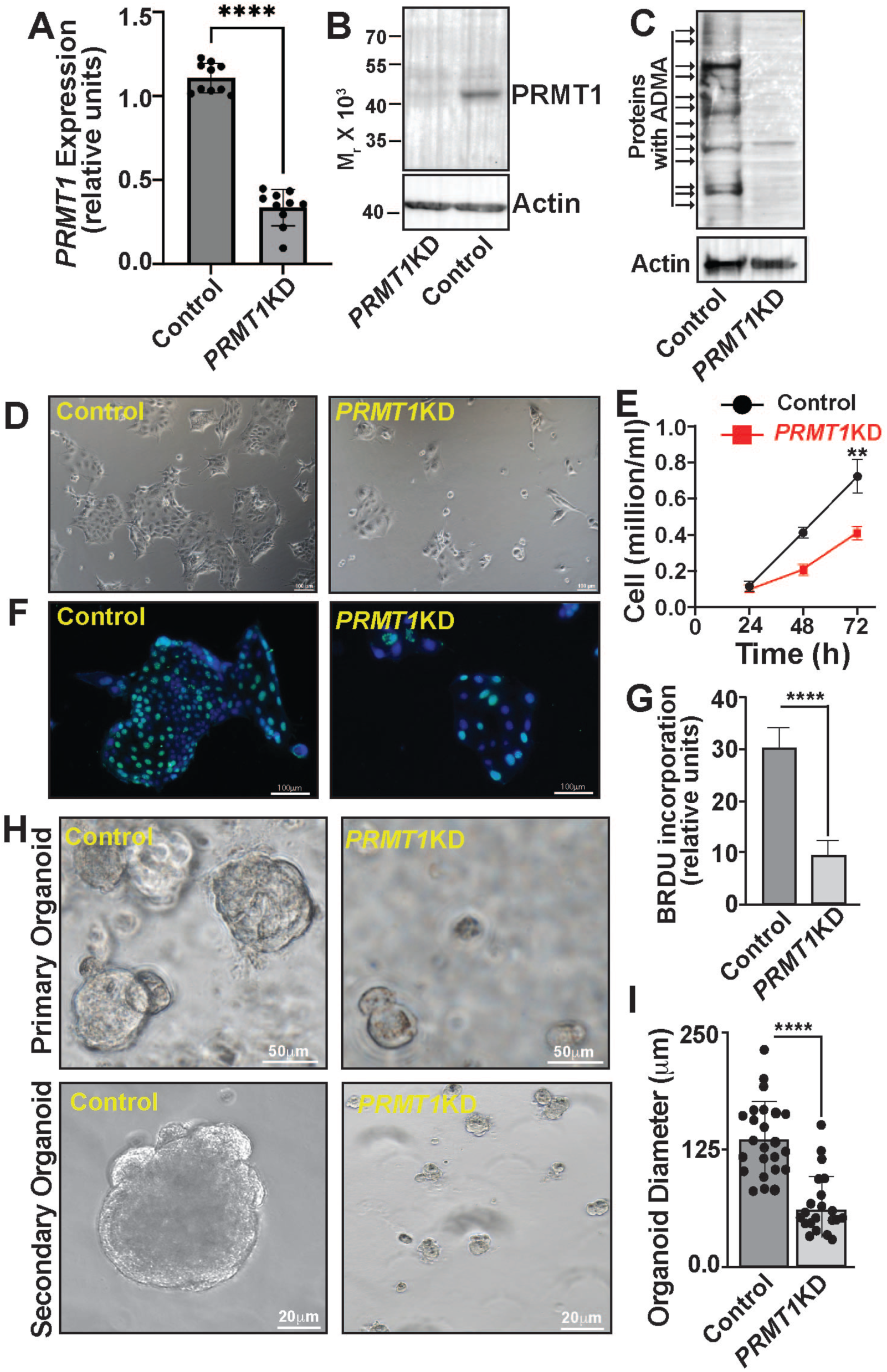
Loss of PRMT1 in hTSCs impairs self-renewal. **(A)** RT-qPCR (n = 3 independent experiments, p≤0.001) and **(B)** Representative western blots, respectively, confirming depletion of PRMT1 expression in *PRMT1-*KD hTSCs. **(C)** Representative western blots confirming loss of Global ADMA in *PRMT1-*KD hTSCs. **(D)** Representative micrographs showing colony morphology of control (CT27) and *PRMT1-KD* hTSCs after 72h in culture. Note smaller colony sizes of the *PRMT1-KD* hTSCs. **(E)** Graphs show reduced cell proliferation in *PRMT1*-KD hTSCs (n = 4 independent experiments, p≤0.005). **(F)** Representative immunostained colonies showing BrdU incorporation in control and *PRMT1-KD* hTSC when cultured in stem-state culture condition for 72 hours. **(G)** BrdU positive nuclei quantitation showing loss of *PRMT1* significantly reduces rate of hTSC proliferation (n=3 independent experiments, p<0.001). **(H)** Micrographs show inefficient organoid formation by *PRMT1*-KD hTSCs. **(I)** Quantitation of primary organoid diameters formed with control and *PRMT1*-KD hTSCs.

We also tested hTSC self-renewal by assessing their ability to form 3-dimensional trophoblast organoids (hTSC organoids)^12,41,42^. Unlike the control hTSCs, *PRMT1-KD* hTSCs showed severe impairment in hTSC organoid formation (Fig. 4H). Control hTSCs formed large organoids, which could be dissociated to single cells and passaged to form secondary organoids, confirming the self-renewing ability (Fig. 4H). In contrast, *PRMT1-KD* hTSCs either did not organize into an organoid structure or formed much smaller organoids and failed to develop secondary organoids upon passaging (Fig. 4H, I), indicating a major defect in self-renewing ability.

### PRMT1 ensures transcriptome fidelity to safeguard premature STB development and is essential for EVT development

We reasoned that the defective self-renewal in *PRMT1*-KD hTSCs should be evident from alteration in TSC stem-state gene expression at single-cell resolution. Thus, we performed scRNA-seq with control and *PRMT1-KD* hTSCs, when maintained in stem-state culture. Seurat cell clustering of scRNA-seq data resulted in 7 cell clusters (Fig. 5A). Majority of control hTSCs clustered to cell clusters 1, 2 and 3, with enrichment of stem-state genes such as, *TEAD4, TP63* and *BCAM*^12,43–45^ (Fig. 5B, C). *PRMT1* expression was also enriched in cells of these undifferentiated TSC Clusters (5B). In contrast, *PRMT1-KD* hTSCs mostly clustered to cell clusters 0, 4, and 5, with reduced expressions of TSC stem–state genes but induced expressions of STB genes, such as *CGA, CGB3* and *KISS1* (Fig. 5B, C). Cells in clusters 0 and 4, also showed enrichment of *ITGA2* and *NOTCH1*, which are induced in cCTBs of a developing human placenta^46,47^ (Fig. 5B). Thus, unbiased scRNA-seq confirmed a major shift in the global gene expression program in *PRMT1-KD* hTSCs with loss of expressions of TSC-stem state genes in majority of cells.

**Figure 5.**
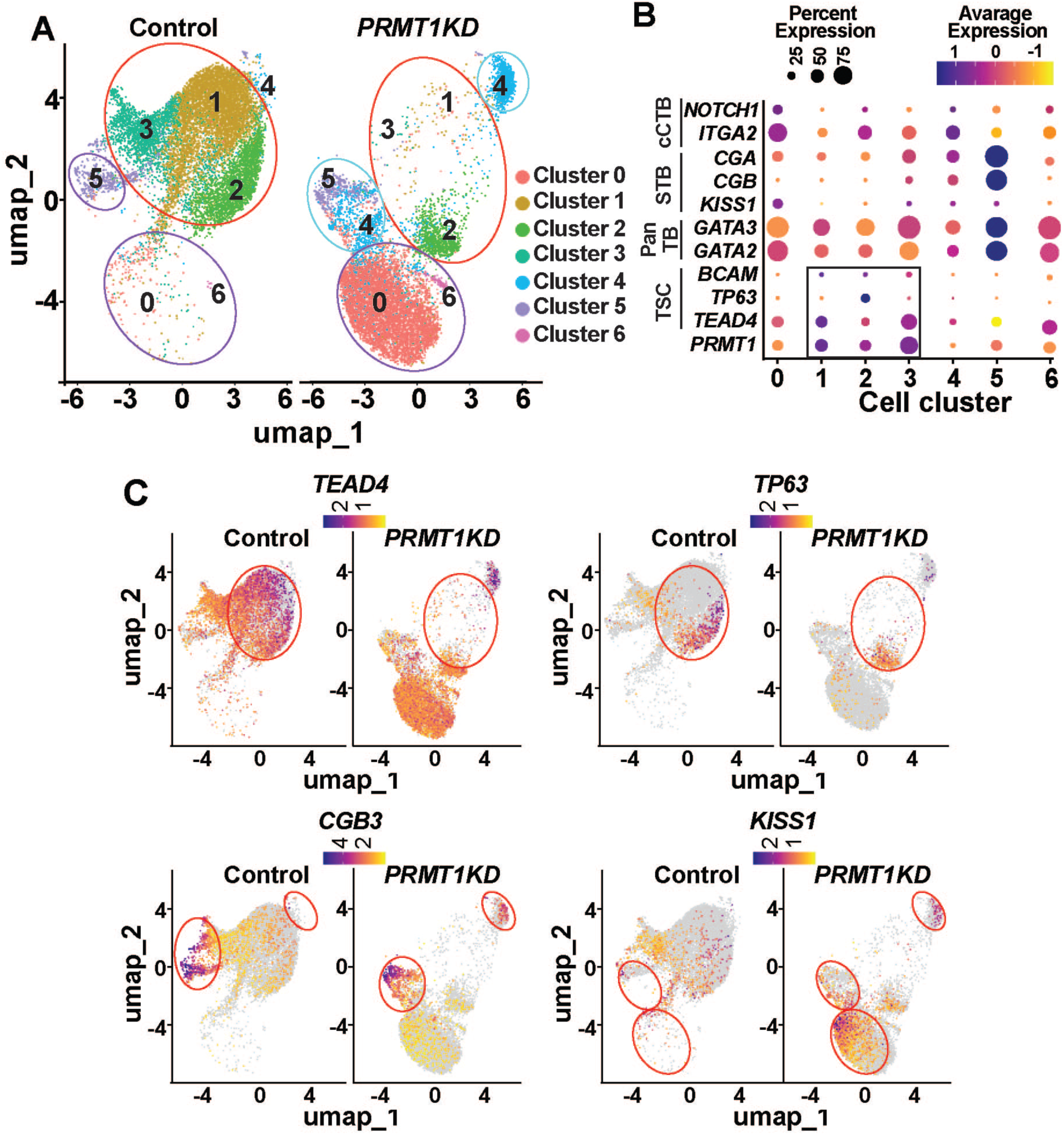
PRMT1 maintains transcriptome fidelity in hTSCs. **(A)** UMAP clustering of the scRNA-Seq data from control and *PRMT1-KD* hTSCs. Clusters were generated according to transcriptome similarity. Note near complete loss of cells in clusters 1,2 and 3 and gain in cell numbers in clusters 0, 4, 5 and 6 for *PRMT1-KD* hTSCs. **(B)** Dot plot showing average expression and percent of cells in each cluster expressing PRMT1 along with pan trophoblast (Pan TB) markers (GATA2, GATA3), hTSC stem state markers (TEAD4, TP63, BCAM), STB markers (CGA, CGB and KISS1) and cCTB markers (ITGA2 and NOTCH1) and novel marker genes identified for each cluster. Genes listed on the x-axis and clusters on the y-axis. **(C)** Single cell feature plots showing expressions of trophoblast marker genes in specific clusters, the scale represents the level of expressions.

To understand how PRMT1 shapes the global gene expression program during human trophoblast development, we also performed bulk RNA-seq analyses in control and *PRMT1-KD* hTSCs. RNA-seq analysis showed significant (1.5 fold) upregulation of 2651 genes and downregulation of 2214 genes in *PRMT1-KD* hTSCs (Fig. 6A, Dataset S2). The RNA-seq analysis also confirmed downregulation of key stem-state regulatory genes (Fig. 6A, Fig. S5A), including *TEAD4* and *MYBL2*, which are not only essential to maintain self-renewal in CTB progenitors but loss of their expressions has been implicated as molecular causes for RPL^12,13^. Quantitative RT-PCR (Fig. S5B) and immunofluorescence analyses (Fig. 6B) also confirmed loss of TEAD4 and MYBL2 expressions in *PRMT1-KD* hTSCs. We also noticed that genes like *NOTUM, MMP11, PLAC8, IFI27* and *DLC1*, which are induced in invasive EVT lineage or are implicated in EVT development^40,48–50^, were also repressed in *PRMT1-KD* hTSCs (Fig. 6A, Fig. S5A). In contrast, genes that are associated with STB development, such as *GCM1*^49,51^*, OVOL1*^52^ along with *CGA, CGB, ERVW-1, PRKCZ and PSG* genes were induced (Fig. 6A, Fig. S5A, S5B). Thus, the unbiased scRNA-seq and bulk RNA-seq analyses in hTSCs indicated that PRMT1 is essential to induce transcription of key stem state and EVT regulators and to prevent induction of STB regulators in trophoblast progenitors.

**Figure 6.**
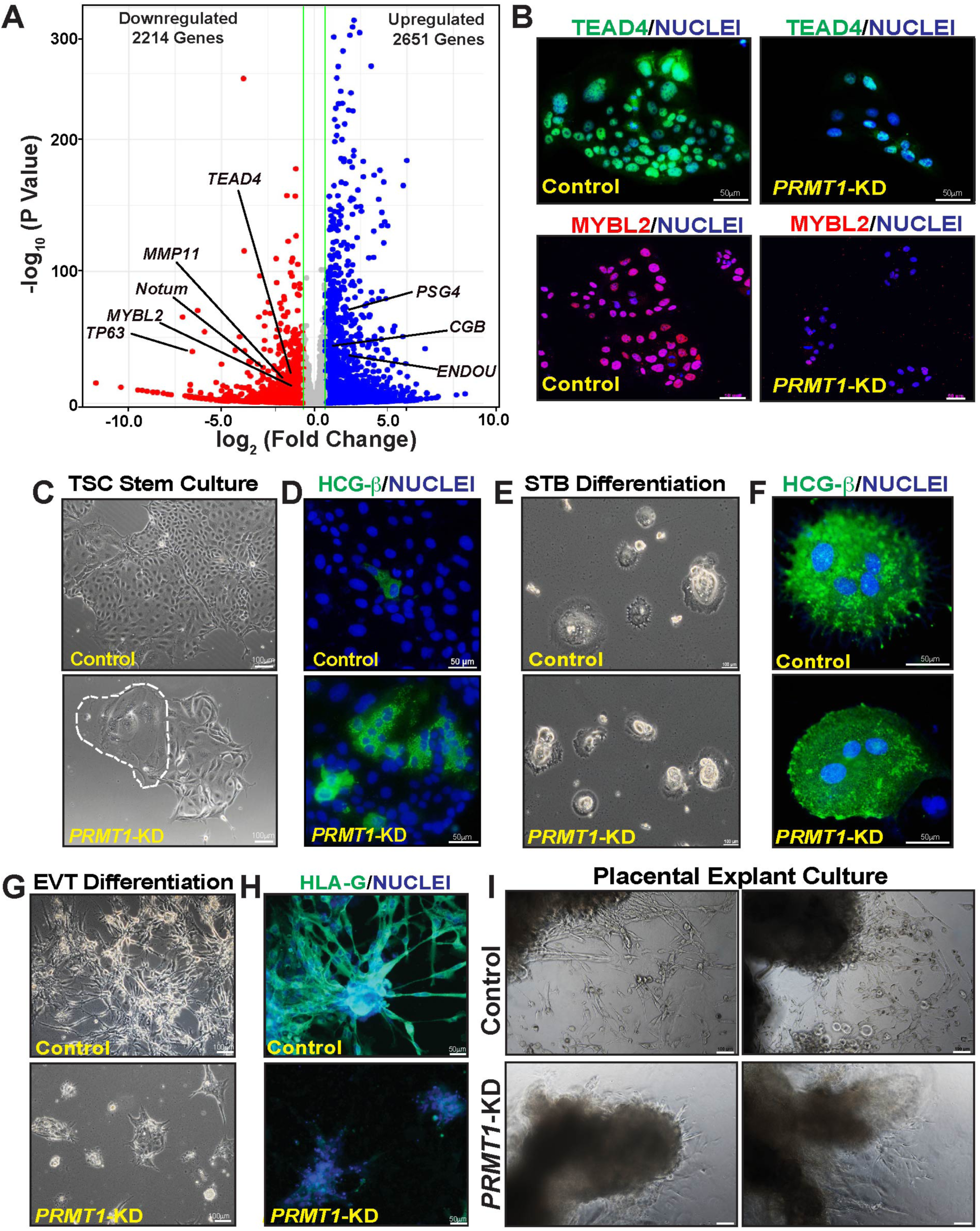
PRMT1 function in hTSCs prevents spontaneous STB development and promotes EVT development. (A) Volcano plot showing global gene expression changes in *PRMT1-KD* hTSCs. Unbiased RNA-seq analyses were performed and ≥1.5-fold gene-expression changes in *PRMT1-KD* hTSCs with a false discovery rate of P < 0.05 were indicated with colored dots (blue: up-regulated, red: down-regulated). Downregulation of *TEAD4 (*a marker of undifferentiated CTBs*), MYBL2 (*important for CTB and EVT development*), NOTUM and MMP11 (*EVT-specific genes*) and* Upregulation of *CGB, PSG4* and *ENDOU* (STB-specific genes) are indicated. (B) Immunostaining images confirming loss of TEAD4 and MYBL2 protein expression in *PRMT1-KD* hTSCs. (C) Representative micrographs showing colony morphology of control and *PRMT1-KD* hTSCs after they were cultured for two passages in stem state culture condition. Note presence of cells with differentiated morphology (white dotted area) in the *PRMT1-KD* hTSC colony. (D) Representative hTSC colonies, maintained at stem state culture condition, showing hCGβ induction upon loss of PRMT1. (E) Representative micrographs showing STB differentiation of control and *PRMT1-KD* hTSCs after they were cultured in STB culture condition. (F) Immunostaining images confirming strong hCGβ induction in both control and *PRMT1-KD* hTSCs. (G) Representative micrographs showing EVT differentiation of control and *PRMT1-KD* hTSCs showing EVT differentiation was inhibited in *PRMT1-KD* hTSCs. (H) Representative immunostained images of HLA-G (EVT marker) showing inefficient EVT differentiation in *PRMT1-KD* hTSCs. (I) Representative micrographs showing that EVT development is impaired from first-trimester human placental explants, when treated with shRNAs against PRMT1.

As STB-specific genes are induced in *PRMT1-KD* hTSCs, we tested whether they show a spontaneous propensity to differentiate to STBs. Intriguingly, we noticed a spontaneous propensity of PRMT1-KD hTSCs to differentiate to STB-like cells, when cultured for a prolonged period of time in hTSC stem-state culture condition. The differentiation to STB-like cells was evident from loss of stem-state colony morphology and induction of STB-specific marker, hCG-β (Fig. 6C, D, Fig. S5C). However, in STB differentiating culture condition, PRMT1-KD hTSCs readily formed multinucleated STBs (Fig. 6E, F). These results indicated that PRMT1 is dispensable for STB development; rather it prevents premature development of STB fate in undifferentiated CTBs/TSCs.

Next, we tested EVT differentiation potential in PRMT1-KD hTSCs by culturing cells in EVT differentiation culture conditions. Loss of PRMT1 in hTSCs strongly inhibited the efficiency of EVT differentiation (Fig. 6G, H, Fig. S5D). To further confirm importance of PRMT1 in EVT development, we tested EVT emergence from human first-trimester placental explants after depleting PRMT1 expression via RNAi. Similar to our findings with hTSCs, EVT emergence from first-trimester placental explants was strongly inhibited upon depletion of PRMT1 expression (Fig. 6I). Collectively, our findings indicated that, during human placentation, PRMT1 balances trophoblast differentiation. It prevents premature STB development but plays an essential role for EVT development.

### Defect in PRMT1 expression inhibits self-renewal ability and EVT development potential in RPL patient-specific hTSCs

To understand the correlation of defective PRMT1 expression in CTB progenitors and EVT precursors with adverse human pregnancies, we established patient-specific hTSC lines from CTBs isolated from placentae associated with idiopathic RPL (RPL-hTSCs). To that end we established hTSC lines from RPL-placentae, in which we noted strong loss of PRMT1 expression in CTBs. These RPL-hTSC lines maintained expression of trophoblast marker GATA3 and micro RNAs from the Chromosome 19 micro-RNA cluster (Fig. S6A, B), which are characteristic markers of hTSCs.

Similar to the CTBs from which they were derived, established RPL-hTSC lines maintained low level of PRMT1 expression (PRMT1_LOW_ RPL-hTSCs) (Fig. 7A, Fig. S7A). The PRMT1_LOW_ RPL-hTSCs showed major defect in cell proliferation (Fig. 7B, Fig. S7B and C) and organoid formation ability (Fig. 7C, D). We also noticed impairment in EVT development potential in PRMT1_LOW_ RPL-hTSCs (Fig. 7E, Fig. S7D). Remarkably, ectopic rescue of PRMT1 expression (Fig. 7F, Fig. S7E) in PRMT1_LOW_ RPL-hTSCs significantly increased cell proliferation (Fig. 7G, Fig. S7F) and EVT differentiation ability (Fig. 7H).

**Figure 7.**
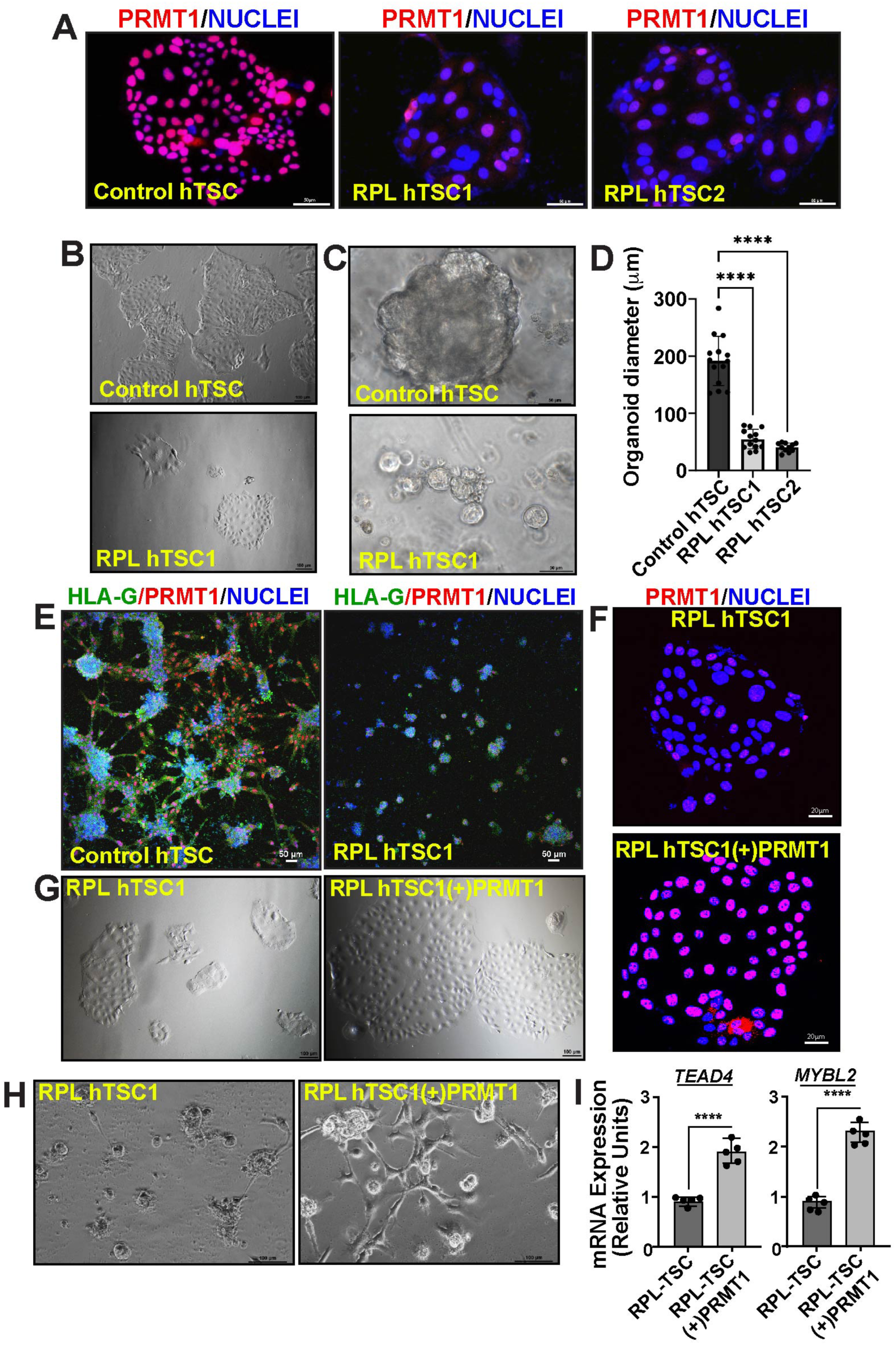
Dysregulated PRMT1 expression in RPL patient-derived hTSCs (RPL-hTSCs) is associated with defective self-renewal ability and inefficient EVT development. (A) Representative immunofluorescence images depicting reduced PRMT1 protein expression in RPL-hTSCs. CT27 hTSC line was used as control. (B) Representative micrographs showing colony morphology of control and one of the RPL-TSC lines, when cultured in stem state culture condition. (C) Micrographs show organoid formation by control CT27 hTSCs and RPL-hTSCs. (D) Quantitation of organoid diameters developed from control CT27 and two RPL-TSC lines. (E) Immunofluorescence images showing defective EVT development and HLA-G induction in RPL-TSCs. (F) Representative immunofluorescence images showing rescue of PRMT1 protein expression in RPL-hTSCs. (G) Representative colony morphology of RPL-TSCs with or without the rescue of PRMT1 expression. Equal number of cells were plated and cultured for 72h. (H) Rescue of PRMT1 expression promotes EVT differentiation in RPL-hTSC lines. Representative micrographs show cell morphology after 6 days of culture in the EVT differentiation condition. (I) RT-qPCR plots show induction of TEAD4 and MYBL2 mRNA expression in RPL-TSC lines upon rescue of PRMT1 expression (n=5 independent experiments, p≤0.01).

Loss of TEAD4 and MYBL2 are implicated in idiopathic RPL and their expressions were down regulated in *PRMT1-*KD hTSCs. Therefore, we tested their expressions in PRMT1_LOW_ RPL-hTSCs. We found that mRNA expression of both *TEAD4* and *MYBL2* are significantly reduced in PRMT1_LOW_ RPL-hTSCs (Fig. S7G) and ectopic rescue of PRMT1 significantly induced expression of both *TEAD4* and *MYBL2* (Fig. 7I). These results indicated that PRMT1 is an upstream regulator of *TEAD4* and *MYBL2* in human trophoblast progenitors and defective TEAD4 and MYBL2 expression due to a defect in PRMT1 expression may be a molecular cause leading to defective trophoblast development and early pregnancy loss.

### A PRMT1-H4R3Me2a epigenetic axis promotes transcription of TEAD4 and MYBL2 in trophoblast progenitors

We asked how loss of PRMT1 in hTSCs reduces transcription of key genes such as TEAD4 and MYBL2. Earlier studies showed that PRMT1-mediated H4R3Me2a modification at chromatin loci establish and maintain transcriptionally active chromatin^28,31^. We noticed that the H4R3Me2a modification is abundantly enriched in the CTBs and EVTs of a first-trimester human placenta and in hTSCs (Fig, 8A,B, C and Fig. S8A). Furthermore, the global H4R3Me2a modification was strongly reduced in PRMT1-KD-hTSCs (Fig. 8B) as well as in PRMT1_LOW_ RPL-hTSCs (Fig. 8C,D). Thus, we hypothesized that PRMT1-mediated histone H4R3Me2a is an epigenetic mechanisms to promote transcription at the chromatin loci of key trophoblast genes such as *TEAD4* and *MYBL2*.

**Figure 8.**
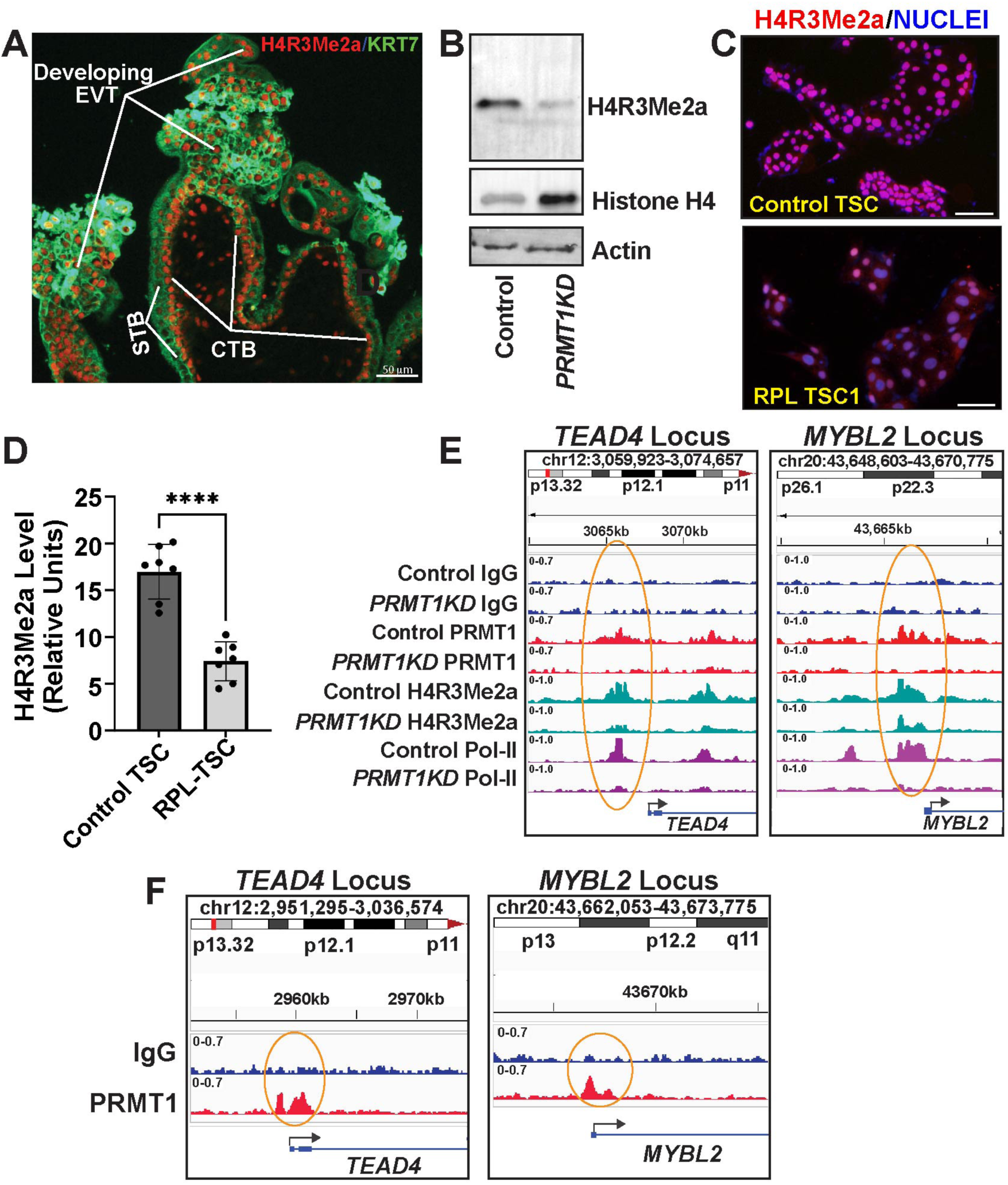
A PRMT1-H4R3Me2a epigenetic axis promotes transcription of key regulators in human trophoblast progenitors. (A) Immunofluorescence analysis showing high level of H4R3Me2a mark in CTBs and EVT precursors in a first-trimester (week 6+4) human placenta. (B) Representative western blots showing loss of H4R3Me2a modification in *PRMT1-KD* hTSCs. Total Histone H4 and β-Actin levels were tested for loading control. (C) Immunofluorescence images showing loss of H4R3Me2a modification in an RPL-TSC line with respect to the CT27 control hTSCs. (D) Image J quantitation of H4R3Me2a modification in control and RPL TSCs. (n=7 independent experiments, performed with 4 different RPL-TSC lines. P≤0.01) (E) Integrative Genome Viewer (IGV) tracks showcasing PRMT1, H4R3Me2a and RNA POL-II CUT&RUN Seq peaks near the *TEAD4* and *MYBL2* promoter regions in control and *PRMT1-KD* hTSCs. Circles indicate statistically significant PRMT1 peaks (red), H4R3Me2a peaks (green) and RNA POL-II peaks (violet) in control TSCs, which are reduced in *PRMT1-KD* hTSCs. (E) IGV tracks showcasing PRMT1 CUT&RUN Seq peaks near the *TEAD4* and *MYBL2* promoter regions in primary CTBs isolated from first-trimester human placentae.

To test our hypothesis, we performed unbiased CUT & RUN-seq and asked three different aspects: (i) PRMT1 recruitment, (ii) H4R3Me2a enrichment and (iii) RNA-polymerase II (POL-II) recruitment at the chromatin loci in control and PRMT1-KD-hTSCs (Fig. 8E). The CUT & RUN-seq identified 1547 PRMT1 target genes in hTSCs including *TEAD4* and *MYBL2* (SI Appendix Data Set S3). Analyses of PRMT1 binding regions revealed that it mainly occupies the promoter regions in trophoblast genome (Fig. S8B) including the promoter regions of *TEAD4* and *MYBL2* (Fig. 8E). HOMER analyses showed that motifs of TEAD and STAT transcription factors are most significantly enriched in PRMT1 binding regions (Fig. S8C). To further confirm that the PRMT1 recruitment at the TEAD4 and MYBL2 loci is an event in primary CTB progenitors of a developing human placenta, we performed CUT&RUN analyses in human CTBs isolated from first-trimester placentae. The CUT&RUN analyses confirmed PRMT1 binding near the *TEAD4* and *MYBL2* promoter region in first-trimester human CTBs (Fig. 8F) The CUT&RUN analyses also confirmed H4R3Me2a enrichment and strong POL-II recruitment at the PRMT1 target loci including the TEAD4 and MYBL2 promoter regions in hTSCs (Fig. 8E). Intriguingly, RNAi mediated depletion of PRMT1 not only reduced PRMT1 recruitment at the *TEAD4* and *MYBL2* promoter regions but also strongly reduced H4R3Me2a enrichment and POL-II recruitment (Fig. 8E), indicating that PRMT1 is required for H4R3Me2a enrichment and POL-II recruitment at these loci.

We also tested PRMT1 recruitment and H4R3Me2a enrichment at the *Tead4* and *Mybl2* loci in the TSPCs of a developing mouse placenta primordium. We isolated ExE/EPC region from E7.0 mouse conceptuses and performed CUT&RUN analyses. Remarkably we noticed that both *Tead4* and *Mybl2* loci are PRMT1 targets in mouse TSPCs (Fig. S8D). We also noticed enrichment of H4R3Me2a modification at PRMT1 binding sites (Fig. S8D). Thus, our unbiased CUT & RUN analyses strongly indicated a conserved PRMT1-H4R3Me2a epigenetic axis that regulates transcription of TEAD4 and MYBL2 in trophoblast progenitors.

## Discussion

Progression of early human pregnancy relies on the fine-tuning of transcriptional programs in CTB progenitors, which balance their self-renewal vs. differentiation to STB and EVTs. Our findings in this study provide strong evidence that PRMT1 is a central player in balancing that gene expression program in human CTB progenitors. We have also identified a conserved epigenetic mechanism, in which a PRMT1-H4R3Me2a regulatory axis shapes the gene expression program in human and mouse trophoblast progenitors. Our study also implicates the importance of this axis to sustain early human pregnancy as dysregulation of this axis is often associated with idiopathic RPL. Based on our observations, we propose that loss of PRMT1 results in loss of H4R3Me2a modification and POL-II recruitment at chromatin loci of key trophoblast genes, leading to defective gene expression, which in turn affects trophoblast development and placentation. To our understanding this is the first report implicating a defect in histone arginine methylation in early pregnancy failure.

Interestingly, the Pawlak et al., study^16^ with global *Prmt1-*KO mice (using a gene-trap allele) predicted that extraembryonic tissues develop normally in homozygous embryos recovered at E6.5. However, the study didn’t include any experiments regarding TSPCs development. Our study in *Prmt1-*cKO mice established that PRMT1 function in TSPCs is essential for progression of pregnancy. However PRMT1 is repressed in developing STBs within the labyrinth zone during mouse placentation (shown in Fig. S2A). Similarly, loss of PRMT1 affects hTSC self-renewal but not STB development. Thus, the essential role of PRMT1 in self-renewing trophoblast progenitors and its dispensability during STB development is a conserved phenomenon. Loss of PRMT1 in hTSCs also inhibits EVT development. However, an earlier study^53^ that conditionally deleted PRMT1 in junctional zone trophoblast cells indicated that the PRMT1 function in mouse junctional zone trophoblasts is also dispensable for placenta and embryonic development. In contrast to human and rat placentation, which are associated with extensive trophoblast invasion^54^ and uterine artery remodeling; the trophoblast invasion during mouse placentation is shallow. Thus, mouse is not an optimum model to study invasive trophoblast development. Given the recent success in invasive trophoblast-specific conditional gene knockout in rat^55^, a future study, using rat as a model system, might be more insightful to test the importance of PRMT1 in invasive trophoblast development *in vivo*.

In a normal developing human placenta PRMT1 is highly expressed in CTB progenitors, and EVT precursors but is suppressed in STBs and idiopathic RPL is often associated with loss of PRMT1 in CTBs and EVT precursors. The differential expression pattern of PRMT1 during human placenta development raises three important questions: (i) Which cellular mechanism induces and maintains PRMT1 expression in undifferentiated CTBs and EVT precursors? (ii) Why is PRMT1 repressed in STBs during normal development and (iii) What leads to defective PRMT1 expression in CTBs and EVT precursors of idiopathic RPL placentae? Distinct signaling mechanisms, such as epidermal growth factor (EGF) and WNT signalling pathways has been implicated to maintain self-renewal in CTBs and CTB-derived hTSCs^40,56,57^, whereas TGFβ signalling has been implicated in EVT development^58^. Thus, it is possible that these signalling pathways are required to promote PRMT1 expression in CTB progenitors and EVT precursors and suppression of these signalling pathways lead to EVT repression in idiopathic RPL CTBs/EVT precursors. However, other possibilities also exist. Considering the rate of idiopathic RPL increases with maternal aging^59,60^, loss of PRMT1 expression in idiopathic RPLs could be acquired mutation at the regulatory *cis*-element, leading to impaired *PRMT1* gene transcription. Our findings in this study will provide an excellent platform for future studies to test mechanisms that are involved in PRMT1 regulation in trophoblast cells during early human placentation.

PRMT1 regulates ADMA modification on many proteins including Histone H4. In this study, we focused on H4R3Me2a modification as we tried to decipher the mechanistic link between PRMT1 and transcriptional regulation of key trophoblast genes like TEAD4 and MYBL2. Our study indicated that along with transcriptional induction of stem state genes, PRMT1 maintains hTSC/CTB stemness by repressing transcriptions of STB-associated genes, such as*, CGBs, PSGs* and *SDC1*. Along with the CTB and STB genes, several hundred other genes were also downregulated or upregulated in *PRMT1-KD* hTSCs. This raises the question about the PRMT1-dependent mechanism that instigates gene induction vs. gene repression. Studies in other cellular contexts have shown that along with H4R3Me2a modification various other PRMT1-mediated mechanisms contribute to gene regulation. PRMT1 could induce gene activation via methylation and stabilization of MLL2^61^. In contrast, PRMT1 could repress genes by altering transcriptional activity of SP1/3^62^. PRMT1 also represses gene expression by inhibiting STAT1 transcriptional activity through methylation of STAT1 inhibitor PIAS1 (protein inhibitor of activated STAT1)^63^. Thus, it is possible that the PRMT1-mediated arginine modification on other proteins indirectly regulates gene expression during human trophoblast development.

Our CUT&RUN study has identified many PRMT1 target genes in hTSCs/CTBs. However, PRMT1 does not have any defined DNA-binding domain. This raises the possibility that PRMT1 binding to trophoblast chromatin loci is mediated along with other transcription factors. Direct interactions of PRMT1 with orphan nuclear receptor HNF4 and FOXO1 have been implicated in recruiting PRMT1 at HNF4 and FOXO1 target promoters^64,65^.However, HNF4 is not expressed in CTBs/hTSCs and FOXO1 expression is extremely low in CTBs/hTSCs^40^. It was also shown that transcription factor USF1 facilitates PRMT1 recruitment at the insulator element of the beta-globin locus^66^. Although USF1 is expressed in CTBs/hTSCs, HOMER analyses of PRMT1 target region indicated no significant enrichment of USF1 binding motif. Rather, the PRMT1 target regions have prevalence of TEAD and STAT motifs. Our mechanistic analyses in this study indicate that PRMT1 is an upstream regulator of TEAD4. However, other TEAD family members, such as TEAD1 and TEAD3 are abundantly expressed in CTBs and EVTs and are implicated in CTB self-renewal^67^ and EVT development^68^. Similarly, various STAT family transcription factors are also expressed in trophoblast cells and implicated in trophoblast biology. For example, STAT1^69^ and STAT3^70^ have been shown to regulate trophoblast proliferation and invasion. Thus, it is possible that certain TEAD and STAT family members facilitate PRMT1 recruitment at target genes. As the majority of PRMT1 binding regions are promoter proximal, it will be interesting to further pursue the interaction of PRMT1 with other transcription factors and how those interactions are involved in trophoblast gene regulation.

The PRMT family constitutes of nine family members and a survey of gene expression revealed that PRMT1 is the most abundantly expressed PRMT family member in CTBs. Furthermore, our studies in *PRMT1-KD* hTSCs indicate that it is the major regulator of ADMA in human trophoblast cells. However, other PRMT family members, except PRMT8, are expressed in CTBs^40^ and may be involved in different aspects of human trophoblast development and function. Thus, our study indicates a new mechanistic paradigm in which PRMT1 and other PRMT family members regulate human trophoblast development and placentation by controlling protein arginine methylation.

## Methods

### Collection and Analyses of Mouse Embryos

All procedures were performed after obtaining IACUC approvals at the University of Kansas Medical Center (KUMC). The *Prmt1*-Tm1a mice (*Prmt1 ^tm1a^*^(EUCOMM)Wtsi^) were obtained from the European Conditional Mouse Mutagenesis Program (EUCOMM). The heterozygous animals were bred to obtain litter and embryos were harvested at different gestational days. Pregnant female animals were identified by presence of vaginal plug at gestational day 0.5, and embryos were harvested at various gestational days. Genomic DNA preparation was done using Extract-N-Amp tissue PCR kit (Sigma, XNAT).

### Generation of PRMT1 Conditional Knockout (cKO) Mice Strains

To generate a conditional allele of PRMT1 (*PRMT1^Flox^*), we employed the Cre/loxP recombination system, targeting exons 4 and 5 of the *PRMT1* gene, which were flanked by loxP sites. Initially, *PRMT1^tm1a^* mice were crossed with *FLP1* recombinase-expressing mice (Jackson Laboratory Strain:009086) to excise the β-galactosidase/neomycin resistance (β-gal/neo) selection cassette via FLP1-mediated recombination at FRT sites. The resulting *PRMT1^Flox/+^* mice were intercrossed to generate homozygous *PRMT1^Flox/Flox^* animals. *PRMT1^Fl/Fl^* mice were bred with *Eomes^CreER^* mice to induce trophoblast progenitor cell-specific deletion of *PRMT1*. *PRMT1^Flox/+;^ Eomes^CreER^* offspring were then crossed with *PRMT1^Flox/Flox^* mice to establish a stable line of*; EomesCreER* animals. For experimental induction of Cre activity, *PRMT1^Flox/Flox^* females were mated with *PRMT1^Flox/Flox^; EomesCreER* males. The presence of a vaginal plug was checked at E 0.5, and tamoxifen (0.2 mg/ gm body weight) (Sigma, T5648) was injected intraperitoneally at embryonic day 5.5 and 6.5. Embryos were harvested at different gestational ages and genotyping and cryosectioning were followed according to the published protocols^71,72^. All mice were maintained in 12 h light/dark cycle, with controlled temperature (20-25 C) and humidity (50-70%) with proper food, water and housing provided by the Institutional Animal Care and Use Committee at KUMC. All genotyping primers are mentioned in Supplementary Table 2.

### Mouse EPC Explant Culture

*PRMT1^tm1a^* male and female mice were crossed, and pregnant female mice were sacrificed at E7.5. Uterine horns were immediately dissected in sterile cold PBS containing 10% FBS (Gibco, 16000-044). ExE/EPCs regions were carefully dissected, washed twice in 10% FBS in PBS and cultured under proliferative condition for 72-96h with MEF-CM medium, which contained 70% MEF-conditioned medium, 30% TS medium, in the presence of 25 ng/ml FGF4 (Invitrogen, PHG0154) and 1 ug/ml Heparin (Sigma, H3149). Explant-outgrowths were measured under microscope and used for further experimental analysis.

### Human trophoblast stem cells culture

Human trophoblast stem cell lines (hTSC) CT27 was gifted by Dr Hiroaki Okae (Tohoku University, Japan). The cells were maintained, passaged, and differentiated as reported in (Okae et al., 2018). The cells were tested for mycoplasma using mycoplasma detection kit (ATCC, 30-1012K).

### Human Placenta samples

Human placental tissues (six to nine weeks of gestation) were obtained from legal pregnancy terminations via the service of Research Centre for Women’s and Infants’ Health (RCWIH) BioBank at Mount Sinai Hospital, Toronto, Ontario, Canada. Placenta Villous samples from various patients with recurrent pregnancy loss (RPL) were obtained after informed consent from all the participants and this study is approved by the Institutional Review Boards at the KUMC. The clinical characteristics of the patients included in this study have been summarized in Supplementary Table S1. Whole placental villi were finely minced, and some portions were fixed in 10% paraformaldehyde for immunohistochemical analysis, while others were rapidly frozen in liquid nitrogen for protein and RNA extraction. The finely dissected villous tissues were used for downstream processing of primary cytotrophoblast cells isolation using immunomagnetic separation using the EasySep™ Human PE Selection Kit (Stemcell Technologies, 17664) and a PE-conjugated anti-ITGA6 (CD49f) antibody (Miltenyi Biotec, 130-127-209), following the manufacturer’s instructions and established protocols^40^. The purified ITGA6-positive cells were seeded onto collagen IV-coated (5 µg/ml) 6-well plates at a density of 0.5–1 × 10**⁶** cells per well and cultured in 2 ml of hTS cell medium.

### Genetic modifications and inhibitors

Lentiviral mediated knockdown was performed in hTS cells using shRNA of PRMT1 (Target sequence: TGAGCGTTCCTAGGCGGTTTC, TRCN0000310243). A scramble shRNA (Addgene-1864, CCTAAGGTTAAGTCGCCCTCGC) was used as control. Lentiviral particles were generated by transfecting plasmids into HEK-293T cells. Virus containing supernatant was collected and virus particles were concentrated by Lenti-X concentrator (Takara, 631231) according to the manufacturer protocol. Human TSCs were transduced using the virous particles at 30-40% confluency. Transduced cells were selected in the presence of puromycin (1-1.5 ug/mL) (Sigma, P8833) and selected cells were checked for knockdown efficiency and used for further experiments.

### Cell Proliferation Assay

Cell proliferation of control scramble and PRMT1 knockdown (*PRMT1*-KD) human trophoblast stem cells (TSCs) were assessed (a) Cell count: 1 × 10**⁵** cells were plated in 6-well culture plates, and cell numbers were recorded at 24-, 48-, and 72-hours post-seeding. Cell growths were monitored using brightfield microscopy (Olympus IX71, Japan). (b) BrdU Assay: 1 × 10**⁴** cells were seeded in 12-well plates, and proliferation was evaluated using the 5-Bromo-2′-deoxyuridine (BrdU) Labeling and Detection Kit I (Roche, Cat. No. 12296736001), following the manufacturer’s protocol. Cells were cultured for 24, 48, and 72 hours, after which nuclei were counterstained with DAPI (BD Biosciences, 564907) and mounted in SlowFade mounting medium (Invitrogen, S36936). BrdU-incorporated nuclei were visualized and quantified under a fluorescence microscope (Nikon Eclipse 80i). Cell death was monitored using Annexin V Apoptosis Detection Kit (Thermo Scientific, 88-8007-72).

### Rescue of PRMT1 in Human TSC line generated from patients with idiopathic RPLs

Lentiviral particles of human PRMT1 (Origene, RC214074L3V) were transduced in Human TSCs at 30-40% confluency. Empty pLKO.1 vector was used as control. Transduced cells were selected in the presence of Puromycin (1-1.5 ug/mL) Selected cells were tested for overexpression efficiency and used for further experiments.

### hTSC organoid generation

Scramble hTSC and PRMT1KD hTSC were used to generate self-renewing organoids as previously described^33^. Briefly, human TSCs were harvested and resuspended in ice-cold trophoblast medium (b-TOM: 10mM HEPES (Sigma H3537), B27(Gibco 17504-044), N2 (Gibco 17502-048) and 2mM glutamine (Gibco 25030081). Growth factor-reduced Matrigel (Corning 354230) was added to reach 60% final concentration. Drops of 25-30 µl containing 40,000 cells were placed in the center of 24-well plates, inverted, and incubated at 37 °C for 20 min to form hanging drops. Plates were then turned upright, and domes were overlaid with 500 µl room-temperature a-TOM (b-TOM plus 100 ng/ml R-spondin, 1 µM A83-01 (PeproTech 120-38), 100 ng/ml rhEGF (Sigma E9644), 50 ng/ml rmHGF (PeproTech 315-23), 2.5 µM Prostaglandin E2 (R&D System, 2296/10), 3 µM CHIR99021 (Sigma, SML1046), and 100 ng/ml Noggin (Invitrogen PHC1506)). Organoids were cultured for 8 days with medium changed every 3 days and appropriate inhibitors added. Brightfield images were taken to monitor growth.

### Placental Explants Culture

Ex vivo placental villous cultures were established as described earlier^34^. Briefly placental villi from 5–8 -week-old gestation placentas were washed in ice-cold 1X PBS and were dissected finely. Pieces containing branching villous-like structures were washed in PBS supplemented with 10% FBS and then subjected to lentiviral particles (scramble and PRMT1 shRNA) for 6 h at 37 °C in a humidified chamber in a 5% CO2/95% air gas mixture. Next, they were encapsulated with growth factor reduced Matrigel (Corning) which was mixed with DMEM/F12 (Gibco 11320-033) on ice to make a Matrigel suspension; 50 µL of the Matrigel suspension was added to each well of a 24-well plate, and explants were placed. The plate was then incubated at 37 °C in a humidified chamber in 5% CO2 for the gel suspension to solidify, thereby encapsulating the explant. Finally, 700 µL EVT (day 1) media was added to each well and allowed culture and media change was done on day 3 and day 6.

### Immunofluorescence

The cells were fixed with 4% paraformaldehyde/PBS at 4°C for 10 minutes, permeabilized with PBS-0.25%TritonX at room temperature for 15 minutes and blocked in PBS-0.1%TritonX-10%FBS at room temperature for an hour. Similarly, the cryosection of human placenta and mouse embryos were also treated in the same way. Then they were incubated with primary antibodies in PBS-0.1%TritonX-10%FBS at 4°C overnight. On the next day, they were stained with secondary antibody in PBS-0.1%TritonX-10%FBS at room temperature for 2 hours. DAPI (BD Biosciences, 564907) counterstain was done and then antifade mounting media (Invitrogen, S36936) were used. Detailed information on antibodies is provided in Supplementary Table S3. For immunohistochemistry, after deparaffinization and rehydration, antigen retrieval with citrate buffer was done, followed by endogenous peroxidase (3% H**₂**O**₂**). DAB counterstaining with Mayer’s Hematoxylin was done and imaged in Nikon Eclipse 80i.

### RNA isolation and RT-qPCR

RNA extraction was carried out using RNeasy mini kit (Qiagen 74104) according to the manufacturer’s protocol. RNA quantification was carried out by Nanodrop (Invitrogen). 1ug of RNA was used for cDNA synthesis using random hexamer primer (Thermo Scientific S0142) and Reverse Transcriptase enzyme (Invitrogen 28025-013) by following standard protocols. qPCR was performed with SYBR Green master mix (Applied Biosystems 4367659) and was performed in biological triplicates and 18sRNA was used as a housekeeping gene for normalization. GraphPad Prism (version) was used for analysis and visualization. Statistical significance was determined by a two-tailed unpaired t-test (****p<0.0001, **p<0.001, **p<0.01, *p<0.05, ns: not 460 significant). Primer sequences for RT-qPCR analyses are provided in Table Supplementary Table S4.

### Protein isolation and western blot

Whole-cell lysates were prepared in RIPA buffer (Thermo Scientific P189900), sonicated, quantified by Bradford assay, boiled in 4x Laemmli buffer (Bio-Rad 1610747), and run on SDS-PAGE (80–100 V). Proteins were transferred to Immunobilon-PVDF membranes on ice at 300 mA for 1 h. Membranes were blocked and incubated with primary antibody (5% skim milk in PBS-T) (BD Biosciences, 232100) overnight at 4 °C, then with secondary antibody at room temperature. Blots were developed using ECL substrate (Sigma WBLURO500) and imaged with a ChemiDOC. Antibody details are in Table Supplementary Table S3 and the uncropped, unprocessed scans are provided in Supplementary Fig. S9.

### CUT&RUN

CUT&RUN experiment using Cell Signaling Technology Kit 86652 was performed according to manufacturer’s instructions. Proliferating 5 × 10^5^ live hTSC cells and primary cytotrophoblasts isolated from 1^st^-Trimester placenta were used per sample for CUT&RUN reactions. In mice, extraembryonic regions including the EPC region were collected at E7.5 from multiple litters to get desired amount of cells. Briefly, cells were captured on Concanavalin A–coated magnetic beads and cell permeabilization was done using buffers containing 0.5% wt/vol Digitonin before incubation with primary antibodies. Protein A and G fused Micrococcal Nuclease was used for DNA digestion, and calcium chloride was added to activate the pAG-MNase enzyme. Stop Buffer was added to each sample and mixed properly, and DNA was purified using Phenol/Chloroform Extraction and Ethanol Precipitation method. The DNA was quantified using 1µL of the freshly dissolved pellet using Qubit instrument (Life Technologies, Waltham, MA). Evaluated the presence of cleaved fragments and the size distribution by capillary electrophoresis with fluorescence detection using a 4200 TapeStation System (Agilent). Samples from three individual CUT&RUN experiments for each experimental condition were used for sequencing. Sequencing libraries were prepared following protocol from Swift bioscience 1S Plus Combinatorial Dual Indexing Kit and Accel-NGS 1S Plus DNA Library Kit. We have used “nf-core-chipseq-2.1.0” pipeline^73^ for downstream analyses of CUT&RUN data. We have used Model-based Analysis of ChIPseq-2 (MACS2) algorithm to call the candidate peaks/binding sites. The MACS2 algorithm uses Poisson distribution to effectively capture local biases and build a model for more robust prediction of peaks. A region was considered to have a significant binding with a p value of <1e-5. The peak distribution in genomic region has been obtained using ChIPseeker v1.42.1. annotatePeak() function and plotted using plotAnotPie() function. Homer 4.11 has been used for annotating the peaks and motif identification.

### Preparation of human trophoblast stem cells for Single Cell-RNA Sequencing

CT27 human trophoblast cells (hTSC) and PRMT1 knockdown hTSC were cultured according to established protocol^40^. The cells were carefully trypsinized and resuspended in hTSC media and cellular viability was checked by automated cell counter (Countess 3, Thermo Fisher Scientific) after staining cells with trypan blue. For the library preparation (done by KUMC genomics core) samples were loaded into the 10X Chromium V3 platform. The raw data for scRNA-seq analyses have been submitted to the Gene Expression Omnibus (GEO) database (https://www.ncbi.nlm.nih.gov/gds), with accession No. GSE295665.

### Preparation of mouse placental cells for Single Cell RNA Sequencing

CD-1 females were paired with CD-1 males, and the presence of a copulation plug was checked daily at various embryonic stages. The day a copulation plug was detected was designated as embryonic day 0.5 (E0.5). Extraembryonic regions, including the ectoplacental cone, were collected at E7.5, and placentae were carefully dissected at E8.5, E9.5, E10.5, E11.5, and E12.5. We ensured to avoid any decidual regions while the dissections. Extraembryonic tissues at E7.5 were obtained from two separate pregnancies due to an insufficient cell yield from a single pregnancy. For stages E8.5 to E12.5, placentae from individual pregnancies were pooled. Tissues (extraembryonic regions or placentae) were digested with collagenase Type IV (30 min at 37 °C). Mechanical dissociation was subsequently performed by serial syringe passage through 18G, 22G, and 25G needles. Cell suspensions were neutralized with 1 mL PBS containing 10% fetal bovine serum (FBS). Released cells were filtered through 40 µm cell strainers and processed using Debris Removal Solution and the Dead Cell Removal Kit (Miltenyi Biotec)). Red blood cells were depleted using an anti-mouse Ter-119 antibody (BD Biosciences, Franklin Lakes, NJ). Cell viability was assessed using an automated cell counter (Countess 3, Thermo Fisher Scientific) following trypan blue staining. Library preparation was performed at the University of Kansas Medical Center (KUMC) Genomics Core Facility, and samples were sequenced using the 10× Genomics Chromium V3 platform. The raw scRNA-seq data have been deposited in the Gene Expression Omnibus (GEO) under accession number GSE278039.

### Single-cell RNA sequencing data analysis

We have used popsicleR v0.2.1 R package to preprocess the single cell RNAseq of different time points (E7.5, E8.5, E9.5, E10.5, E11.5, and E12.5) for mouse Single cell RNA Seq analysis. The low-quality reads and mitochondrial contamination have been removed using Filterplots() function, including the parameters like minimum of molecules detected within a cell being 500 and maximum percentage of mitochondria as 5%. In the next step, the doublets were removed by first identification of cut-off and then removed them using CalculateDoublets() function. SelectIntegrationFeatures of Seurat v4.0.6 R package function screens the features that will be used while performing integration and FindIntegrationAnchors was implemented to get anchors. Finally, IntegrateData function was used to create integrated assay. We used DefaultAssay() function to make this integrated assay default, which will be used in downstream analysis. We used RunPCA function to perform PCA. Next characterization of the principal components and estimate the number of significant PCs that capture the signal of interest, while minimizing noise was accessed using ElbowPlot and DimHeatmap functions. Further functions like FindNeighbors and FindClusters were executed sequentially for clustering. In the final steps dimensional reduction methods UMAP was performed using RunUMAP function of Seurat. Differentially expressed genes were identified using FindAllMarkers()_function and cell clusters were identified based on the expression of marker genes. To further analyze trophoblast cell cluster, data from initial clustering was subsetted. Further it was analyzed using the function subset() function based upon annotations from marker genes identified by FindAllMarkers. Integration of the trophoblast only datasets were performed using the same integration methods above (using 20 dimensions for FindIntegrationAnchors() and IntegrateData(), and 20 PCs and a resolution of 0.5 for FindClusters() and RunUMAP(). The feature plot has been obtained using Featureplot_scCustom() function of scCustomize v3.1.3 R package.

### Single Cell Proportion test

The variability between biological samples used for scRNA-Seq can be high due to variability in the source of the samples, technical reasons like the dissociation protocol used or intrinsic properties of the sample. Permutation_test() is the single cell proportion test function of the R library (https://github.com/rpolicastro/scProportionTest) that analyses scRNA-Seq samples to account for variability in the proportions of cell clusters between samples. A permutation test first calculates a p-value for each cluster, then bootstrapping is used to calculate the confidence interval for the magnitude difference. The result is plotted using permutation_plot() function of scProportion Test R package.

### RNA-Seq analyses

RNA-seq analysis was performed according to published protocol^34^. Total RNA from the control human TSCs and PRMT1-KD TSCs were isolated using RNeasy Mini Kit following manufacturer’s protocol. Integrity of the total RNA samples was evaluated using an Agilent Technologies 2100 Bioanalyzer. The total RNA fraction was processed by oligo dT bead capture of mRNA, fragmentation, and reverse transcription into cDNA. After ligation with the appropriate Unique Dual Index (UDI) adaptors, the cDNA library was prepared using the Universal Plus mRNA-seq +UDI library preparation kit (NuGEN; Marion, SD). Adaptor and low-quality reads were removed using Rfastp R package. Reads were mapped to reference genome hg38 using Rsubread mapping tool. Mapping of reads were done using featureCount tool of subread package. The downstream analysis was done using DEseq2 tool. The raw data for RNA-seq analyses have been submitted to the Gene Expression Omnibus (GEO) database (https://www.ncbi.nlm.nih.gov/gds), with accession No. GSE295904.

### Statistical Analyses

Statistical significance was determined for quantitative RT-PCR analyses for mRNA expression. We performed at least n = 4 experimental replicates for all these experiments. For statistical significance of generated data, statistical comparisons between two means were determined with Student’s *t* test, and significantly altered values (*P* ≤ 0.01) are highlighted in the figures by an asterisk. RNA-seq data were generated with n = 3 experimental replicates per group. The statistical significance of altered gene expression (absolute fold change ≥1.5 and false discovery rate q-value ≤ 0.05) was initially confirmed with right-tailed Fisher’s exact test. Independent datasets were analyzed using GraphPad Prism software.

### GEO Database Accession Codes for Sequencing Data

The raw genomics data is submitted to the GEO database. Data for scRNA-seq of E7.5-E12.5 mouse placentae are submitted with GEO accession number GSE278039. Raw data for bulk RNA-seq, scRNA-Seq in human TSCs and *PRMT1-KD* TSCs and CUT & RUN seq in human TSCs and PRMT1KD TSCs are submitted with GEO accession numbers GSE295904, GSE295665 and GSE295846 respectively. GEO accession number for Bulk RNA-seq for RPL placenta samples is GSE306548.

### Ethics Statement regarding studies with mouse model and human placental tissues

All studies with mouse models were approved by IACUC at the University of Kansas Medical Center (KUMC). Human placental tissues were obtained at the University of Kansas Medical Center (KUMC) and from the Research Centre for Women’s and Infants’ Health (RCWIH) Biobank at the Mount Sinai Hospital, Toronto, Canada. The Institutional Review Boards at the KUMC and at the Mount Sinai hospital approved utilization of human placental tissues and all experimental procedures.

## Author Contributions

Soumen Paul conceived the project, designed experiments and wrote the manuscript. Purbasa Dasgupta designed and performed experiments, analyzed data and wrote initial version of the manuscript. Rajnish Kumar performed all genomic sequence data processing and downstream bioinformatics analyses. Soma Ray worked with hTSCs and worked with animal models. Namrata Roy worked to maintain animal colonies. Asef Jawad Niloy and Mounika Vallakati worked on establishing and culturing RPL-hTSCs. Courtney Marsh provided placental samples. Sebastian Arnold provided the *Eomes*^CreER^ mouse strain and helped with the manuscript ---.

## Acknowledgement

This research was supported by NIH grants HD113673, HD103161, HD062546, HD101319 and HD119510 to S.P. S.J.A. is supported by the German Research Foundation (DFG) through the Heisenberg Program (AR 732/4-1), project grant (AR 732/2-1), project A03 of CRC 850 (project ID 89986987), project A08 of CRC 992 (project ID 192904750) and Germany’s Excellence Strategy (CIBSS – EXC-2189 – Project ID 390939984). We thank Dr. Mark T. Bedford of MD Anderson Cancer Center for providing antibodies for ADMA modifications. This study was also supported by various core facilities, including the Genomics Core, Flowcytometry core, Imaging and Histology Core facility of the University of Kansas Medical Center. We also acknowledge administrative support from the Department of Pathology Internal Medicine and the Institute for Reproduction and Developmental Sciences of the University of Kansas Medical Center.

## References

1 Vasconcelos, S. et al. Immune Dysregulation and Trophoblastic Dysfunction as a Potential Cause of Idiopathic Recurrent Pregnancy Loss. Biology (Basel) 14, doi:10.3390/biology14070811 (2025).

2 Soares, M. J. et al. Regulatory pathways controlling the endovascular invasive trophoblast cell lineage. The Journal of reproduction and development 58, 283–287 (2012).

3 Harris, L. K. Review: Trophoblast-vascular cell interactions in early pregnancy: how to remodel a vessel. Placenta 31 **Suppl**, S93–98, doi:10.1016/j.placenta.2009.12.012 (2010).

4 Kaufmann, P., Black, S. & Huppertz, B. Endovascular trophoblast invasion: implications for the pathogenesis of intrauterine growth retardation and preeclampsia. Biol Reprod 69, 1–7, doi:10.1095/biolreprod.102.014977 (2003).

5 Haider, S. et al. Notch1 controls development of the extravillous trophoblast lineage in the human placenta. Proc Natl Acad Sci U S A 113, E7710–E7719, doi:10.1073/pnas.1612335113 (2016).

6 Beer, A. E. & Sio, J. O. Placenta as an immunological barrier. Biology of reproduction 26, 15–27, doi:10.1095/biolreprod26.1.15 (1982).

7 Costa, M. A. The endocrine function of human placenta: an overview. Reprod Biomed Online 32, 14–43, doi:10.1016/j.rbmo.2015.10.005 (2016).

8 PrabhuDas, M. et al. Immune mechanisms at the maternal-fetal interface: perspectives and challenges. Nat Immunol 16, 328–334, doi:10.1038/ni.3131 (2015).

9 Chamley, L. W. et al. Review: where is the maternofetal interface? Placenta 35 **Suppl**, S74–80, doi:10.1016/j.placenta.2013.10.014 (2014).

10 Knofler, M. et al. Human placenta and trophoblast development: key molecular mechanisms and model systems. Cellular and molecular life sciences: CMLS, doi:10.1007/s00018-019-03104-6 (2019).

11 Boss, A. L., Chamley, L. W. & James, J. L. Placental formation in early pregnancy: how is the centre of the placenta made? Hum Reprod Update 24, 750–760, doi:10.1093/humupd/dmy030 (2018).

12 Saha, B. et al. TEAD4 ensures postimplantation development by promoting trophoblast self-renewal: An implication in early human pregnancy loss. Proceedings of the National Academy of Sciences of the United States of America, doi:10.1073/pnas.2002449117 (2020).

13 Wu, Z. H. et al. Dysregulation of MYBL2 impairs extravillous trophoblast lineage development and function, contributing to recurrent spontaneous abortion. Proceedings of the National Academy of Sciences of the United States of America 122, e2421653122, doi:10.1073/pnas.2421653122 (2025).

14 Jahan, S. & Davie, J. R. Protein arginine methyltransferases (PRMTs): role in chromatin organization. Adv Biol Regul 57, 173–184, doi:10.1016/j.jbior.2014.09.003 (2015).

15 Al-Hamashi, A. A., Diaz, K. & Huang, R. Non-Histone Arginine Methylation by Protein Arginine Methyltransferases. Curr Protein Pept Sci 21, 699–712, doi:10.2174/1389203721666200507091952 (2020).

16 Pawlak, M. R., Scherer, C. A., Chen, J., Roshon, M. J. & Ruley, H. E. Arginine N-methyltransferase 1 is required for early postimplantation mouse development, but cells deficient in the enzyme are viable. Molecular and cellular biology 20, 4859–4869, doi:10.1128/MCB.20.13.4859-4869.2000 (2000).

17 Yu, Z., Chen, T., Hebert, J., Li, E. & Richard, S. A mouse PRMT1 null allele defines an essential role for arginine methylation in genome maintenance and cell proliferation. Molecular and cellular biology 29, 2982–2996, doi:10.1128/MCB.00042-09 (2009).

18 Tee, W. W. et al. Prmt5 is essential for early mouse development and acts in the cytoplasm to maintain ES cell pluripotency. Genes & development 24, 2772–2777, doi:10.1101/gad.606110 (2010).

19 Yoshimoto, T. et al. The arginine methyltransferase PRMT2 binds RB and regulates E2F function. Exp Cell Res 312, 2040–2053, doi:10.1016/j.yexcr.2006.03.001 (2006).

20 Swiercz, R., Cheng, D., Kim, D. & Bedford, M. T. Ribosomal protein rpS2 is hypomethylated in PRMT3-deficient mice. The Journal of biological chemistry 282, 16917–16923, doi:10.1074/jbc.M609778200 (2007).

21 Neault, M., Mallette, F. A., Vogel, G., Michaud-Levesque, J. & Richard, S. Ablation of PRMT6 reveals a role as a negative transcriptional regulator of the p53 tumor suppressor. Nucleic Acids Res 40, 9513–9521, doi:10.1093/nar/gks764 (2012).

22 Kim, J. D. et al. PRMT8 as a phospholipase regulates Purkinje cell dendritic arborization and motor coordination. Sci Adv 1, e1500615, doi:10.1126/sciadv.1500615 (2015).

23 Vikhe, P. P., Tateossian, H., Bharj, G., Brown, S. D. M. & Hood, D. W. Mutation in Fbxo11 Leads to Altered Immune Cell Content in Jeff Mouse Model of Otitis Media. Front Genet 11, 50, doi:10.3389/fgene.2020.00050 (2020).

24 Blanc, R. S., Vogel, G., Chen, T., Crist, C. & Richard, S. PRMT7 Preserves Satellite Cell Regenerative Capacity. Cell Rep 14, 1528–1539, doi:10.1016/j.celrep.2016.01.022 (2016).

25 Yadav, N. et al. Specific protein methylation defects and gene expression perturbations in coactivator-associated arginine methyltransferase 1-deficient mice. Proceedings of the National Academy of Sciences of the United States of America 100, 6464–6468, doi:10.1073/pnas.1232272100 (2003).

26 Yang, Y. & Bedford, M. T. Protein arginine methyltransferases and cancer. Nature reviews. Cancer 13, 37–50, doi:10.1038/nrc3409 (2013).

27 Gayatri, S. & Bedford, M. T. Readers of histone methylarginine marks. Biochim Biophys Acta 1839, 702–710, doi:10.1016/j.bbagrm.2014.02.015 (2014).

28 Thiebaut, C., Eve, L., Poulard, C. & Le Romancer, M. Structure, Activity, and Function of PRMT1. Life (Basel) 11, doi:10.3390/life11111147 (2021).

29 Yang, Y. et al. TDRD3 is an effector molecule for arginine-methylated histone marks. Mol Cell 40, 1016–1023, doi:10.1016/j.molcel.2010.11.024 (2010).

30 Yao, B. et al. PRMT1-mediated H4R3me2a recruits SMARCA4 to promote colorectal cancer progression by enhancing EGFR signaling. Genome Med 13, 58, doi:10.1186/s13073-021-00871-5 (2021).

31 Huang, S., Litt, M. & Felsenfeld, G. Methylation of histone H4 by arginine methyltransferase PRMT1 is essential in vivo for many subsequent histone modifications. Genes & development 19, 1885–1893, doi:10.1101/gad.1333905 (2005).

32 Hemberger, M., Udayashankar, R., Tesar, P., Moore, H. & Burton, G. J. ELF5-enforced transcriptional networks define an epigenetically regulated trophoblast stem cell compartment in the human placenta. Human Molecular Genetics, 1–12 (2010).

33 Home, P. et al. Genetic redundancy of GATA factors in the extraembryonic trophoblast lineage ensures the progression of preimplantation and postimplantation mammalian development. Development 144, 876–888, doi:10.1242/dev.145318 (2017).

34 Ray, S. et al. Hippo signaling cofactor, WWTR1, at the crossroads of human trophoblast progenitor self-renewal and differentiation. Proceedings of the National Academy of Sciences of the United States of America 119, e2204069119, doi:10.1073/pnas.2204069119 (2022).

35 Saha, B. et al. TEAD4 ensures postimplantation development by promoting trophoblast self-renewal: An implication in early human pregnancy loss. Proceedings of the National Academy of Sciences of the United States of America 117, 17864–17875, doi:10.1073/pnas.2002449117 (2020).

36 Austin, C. P. et al. The knockout mouse project. Nature genetics 36, 921–924, doi:10.1038/ng0904-921 (2004).

37 Pimeisl, I. M. et al. Generation and characterization of a tamoxifen-inducible Eomes(CreER) mouse line. Genesis 51, 725–733, doi:10.1002/dvg.22417 (2013).

38 Knofler, M. et al. Human placenta and trophoblast development: key molecular mechanisms and model systems. Cellular and molecular life sciences: CMLS 76, 3479–3496, doi:10.1007/s00018-019-03104-6 (2019).

39 Zhu, J. et al. Whole-Transcriptome Analysis Identifies Gender Dimorphic Expressions of Mrnas and Non-Coding Rnas in Chinese Soft-Shell Turtle (Pelodiscus sinensis). Biology (Basel) 11, doi:10.3390/biology11060834 (2022).

40 Okae, H. et al. Derivation of Human Trophoblast Stem Cells. Cell Stem Cell 22, 50–63 e56, doi:10.1016/j.stem.2017.11.004 (2018).

41 Zhou, J. et al. The immune checkpoint molecule, VTCN1/B7-H4, guides differentiation and suppresses proinflammatory responses and MHC class I expression in an embryonic stem cell-derived model of human trophoblast. Front Endocrinol (Lausanne) 14, 1069395, doi:10.3389/fendo.2023.1069395 (2023).

42 Shannon, M. J. et al. Single-cell assessment of primary and stem cell-derived human trophoblast organoids as placenta-modeling platforms. Developmental cell 59, 776–792 e711, doi:10.1016/j.devcel.2024.01.023 (2024).

43 Li, Y. et al. BMP4-directed trophoblast differentiation of human embryonic stem cells is mediated through a DeltaNp63+ cytotrophoblast stem cell state. Development 140, 3965–3976, doi:10.1242/dev.092155 (2013).

44 Soncin, F., Natale, D. & Parast, M. M. Signaling pathways in mouse and human trophoblast differentiation: a comparative review. Cellular and molecular life sciences: CMLS 72, 1291–1302, doi:10.1007/s00018-014-1794-x (2015).

45 Shannon, M. J. et al. Cell trajectory modeling identifies a primitive trophoblast state defined by BCAM enrichment. Development 149, doi:10.1242/dev.199840 (2022).

46 Lee, C. Q. E. et al. Integrin alpha2 marks a niche of trophoblast progenitor cells in first trimester human placenta. Development 145, doi:10.1242/dev.162305 (2018).

47 Dietrich, B., Haider, S., Meinhardt, G., Pollheimer, J. & Knofler, M. WNT and NOTCH signaling in human trophoblast development and differentiation. Cellular and molecular life sciences: CMLS 79, 292, doi:10.1007/s00018-022-04285-3 (2022).

48 Shukla, V. et al. NOTUM-mediated WNT silencing drives extravillous trophoblast cell lineage development. Proceedings of the National Academy of Sciences of the United States of America 121, e2403003121, doi:10.1073/pnas.2403003121 (2024).

49 Shimizu, T. et al. CRISPR screening in human trophoblast stem cells reveals both shared and distinct aspects of human and mouse placental development. Proceedings of the National Academy of Sciences of the United States of America 120, e2311372120, doi:10.1073/pnas.2311372120 (2023).

50 Chang, W. L. et al. PLAC8, a new marker for human interstitial extravillous trophoblast cells, promotes their invasion and migration. Development 145, doi:10.1242/dev.148932 (2018).

51 Jeyarajah, M. J. et al. The multifaceted role of GCM1 during trophoblast differentiation in the human placenta. Proceedings of the National Academy of Sciences of the United States of America 119, e2203071119, doi:10.1073/pnas.2203071119 (2022).

52 Renaud, S. J. et al. OVO-like 1 regulates progenitor cell fate in human trophoblast development. Proceedings of the National Academy of Sciences of the United States of America 112, E6175–6184, doi:10.1073/pnas.1507397112 (2015).

53 Sato, A. et al. Gestational changes in PRMT1 expression of murine placentas. Placenta 65, 47–54, doi:10.1016/j.placenta.2018.04.001 (2018).

54 Soares, M. J., Chakraborty, D., Karim Rumi, M. A., Konno, T. & Renaud, S. J. Rat placentation: an experimental model for investigating the hemochorial maternal-fetal interface. Placenta 33, 233–243, doi:10.1016/j.placenta.2011.11.026 (2012).

55 Iqbal, K. et al. Conditionally mutant animal model for investigating the invasive trophoblast cell lineage. Development 151, doi:10.1242/dev.202239 (2024).

56 Haider, S. et al. Self-Renewing Trophoblast Organoids Recapitulate the Developmental Program of the Early Human Placenta. Stem cell reports, doi:10.1016/j.stemcr.2018.07.004 (2018).

57 Turco, M. Y. et al. Trophoblast organoids as a model for maternal-fetal interactions during human placentation. Nature 564, 263–267, doi:10.1038/s41586-018-0753-3 (2018).

58 Haider, S. et al. Transforming growth factor-beta signaling governs the differentiation program of extravillous trophoblasts in the developing human placenta. Proceedings of the National Academy of Sciences of the United States of America 119, e2120667119, doi:10.1073/pnas.2120667119 (2022).

59 Li, T. C. et al. An analysis of the pattern of pregnancy loss in women with recurrent miscarriage. Fertil Steril 78, 1100–1106 (2002).

60 Sauer, M. V. Reproduction at an advanced maternal age and maternal health. Fertil Steril 103, 1136–1143, doi:10.1016/j.fertnstert.2015.03.004 (2015).

61 An, D. et al. PRMT1-mediated methylation regulates MLL2 stability and gene expression. Nucleic Acids Res 53, doi:10.1093/nar/gkae1227 (2025).

62 Lafleur, V. N., Richard, S. & Richard, D. E. Transcriptional repression of hypoxia-inducible factor-1 (HIF-1) by the protein arginine methyltransferase PRMT1. Mol Biol Cell 25, 925–935, doi:10.1091/mbc.E13-07-0423 (2014).

63 Weber, S. et al. PRMT1-mediated arginine methylation of PIAS1 regulates STAT1 signaling. Genes & development 23, 118–132, doi:10.1101/gad.489409 (2009).

64 Barrero, M. J. & Malik, S. Two functional modes of a nuclear receptor-recruited arginine methyltransferase in transcriptional activation. Mol Cell 24, 233–243, doi:10.1016/j.molcel.2006.09.020 (2006).

65 Yamagata, K. et al. Arginine methylation of FOXO transcription factors inhibits their phosphorylation by Akt. Mol Cell 32, 221–231, doi:10.1016/j.molcel.2008.09.013 (2008).

66 Huang, S., Li, X., Yusufzai, T. M., Qiu, Y. & Felsenfeld, G. USF1 recruits histone modification complexes and is critical for maintenance of a chromatin barrier. Molecular and cellular biology 27, 7991–8002, doi:10.1128/MCB.01326-07 (2007).

67 Dong, C. et al. A genome-wide CRISPR-Cas9 knockout screen identifies essential and growth-restricting genes in human trophoblast stem cells. Nat Commun 13, 2548, doi:10.1038/s41467-022-30207-9 (2022).

68 Meinhardt, G. et al. The multifaceted roles of the transcriptional coactivator TAZ in extravillous trophoblast development of the human placenta. Proceedings of the National Academy of Sciences of the United States of America 122, e2426385122, doi:10.1073/pnas.2426385122 (2025).

69 Pereira de Sousa, F. L., et al. Involvement of STAT1 in proliferation and invasiveness of trophoblastic cells. Reprod Biol 17, 218–224, doi:10.1016/j.repbio.2017.05.005 (2017).

70 Fitzgerald, J. S., Poehlmann, T. G., Schleussner, E. & Markert, U. R. Trophoblast invasion: the role of intracellular cytokine signalling via signal transducer and activator of transcription 3 (STAT3). Hum Reprod Update 14, 335–344, doi:10.1093/humupd/dmn010 (2008).

71 Bhattacharya, B. et al. Atypical protein kinase C iota (PKClambda/iota) ensures mammalian development by establishing the maternal-fetal exchange interface. Proceedings of the National Academy of Sciences of the United States of America 117, 14280–14291, doi:10.1073/pnas.1920201117 (2020).

72 Ghosh, A. et al. The GATA transcriptional program dictates cell fate equilibrium to establish the maternal-fetal exchange interface and fetal development. Proceedings of the National Academy of Sciences of the United States of America 121, e2310502121, doi:10.1073/pnas.2310502121 (2024).

73 Ewels, P. A. et al. The nf-core framework for community-curated bioinformatics pipelines. Nature biotechnology 38, 276–278, doi:10.1038/s41587-020-0439-x (2020).

