## Supplementary Figures and Tables for "Arginine methyltransferase PRMT1 equipoises trophoblast development to prevent early pregnancy loss"

**Description of Supplementary Data**

**Supplementary Figures S1-S9**

**Supplementary Table 1:** Details of Idiopathic RPL Placental Tissues Used for The Study

**Supplementary Table 2:** Primers Used for Mice Genotyping

**Supplementary Table 3:** Antibodies Used For The Study

**Supplementary Table 4:** Primers used for the RT-qPCR

### Fig. S1

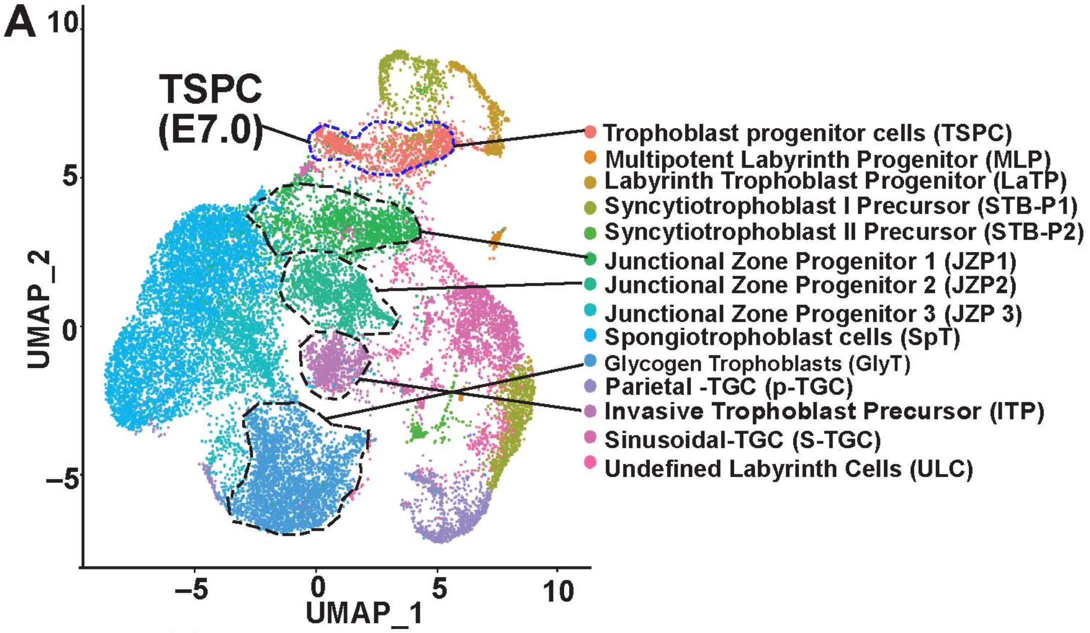


**B**

| Cell Type | Cell Count | Markers |
|---|---|---|
| TSPC | 1254 | *Cdx2, Eomes, Elf5, Esrrb* |
| MLP | 115 | *Epcam, Egfr, Ly6a, Met* |
| LaTP | 425 | *Epcam, Tfrc, Slc16a1, Slc16a3* |
| STB-P1 | 1344 | *Slc16a1,Gjb3, Nr6a1, Dlx3, Syna* |
| STB-P2 | 207 | *Slc16a3, Vegf, Wnt7b* |
| JZP1 | 1953 | *Hand1, Plac1, Rhox9* |
| JZP2 | 1302 | *Hand1, Plac1, Rhox9, Procr, Prl3d1* |
| JZP3 | 1585 | *Hand1, Plac1, Rhox9, Tpbpa* |
| SpT | 5315 | *Cdx2, Eomes, Elf5, Esrrb* |
| GlyT | 2936 | *Cdx2, Ascl2* |
| ITP | 914 | *Prl7b1, Hand1, Prdm1* |
| P-TGC | 861 | *Prl3d1, Prl2c2, Hand1, Procr,* |
| S-TGC | 1544 | *Lifr, Procr, Hand1, Pdgfa, Plac1* |
| ULC | 850 | *Epcam, GJB3* |

**Figure S1**: **(A)** UMAP Clustering of scRNA-Seq data with E7.0-E12.0 mouse placentae showing different trophoblast cell types. Clusters were annotated according to canonical marker genes listed in (B). Cell clusters representative of TSPCs and different junctional zone progenitors, which express high levels of Prmt1 (shown in main Fig. 1A) are marked. **(B)** Marker genes that were used to assign trophoblast cell clusters.

Fig. S2

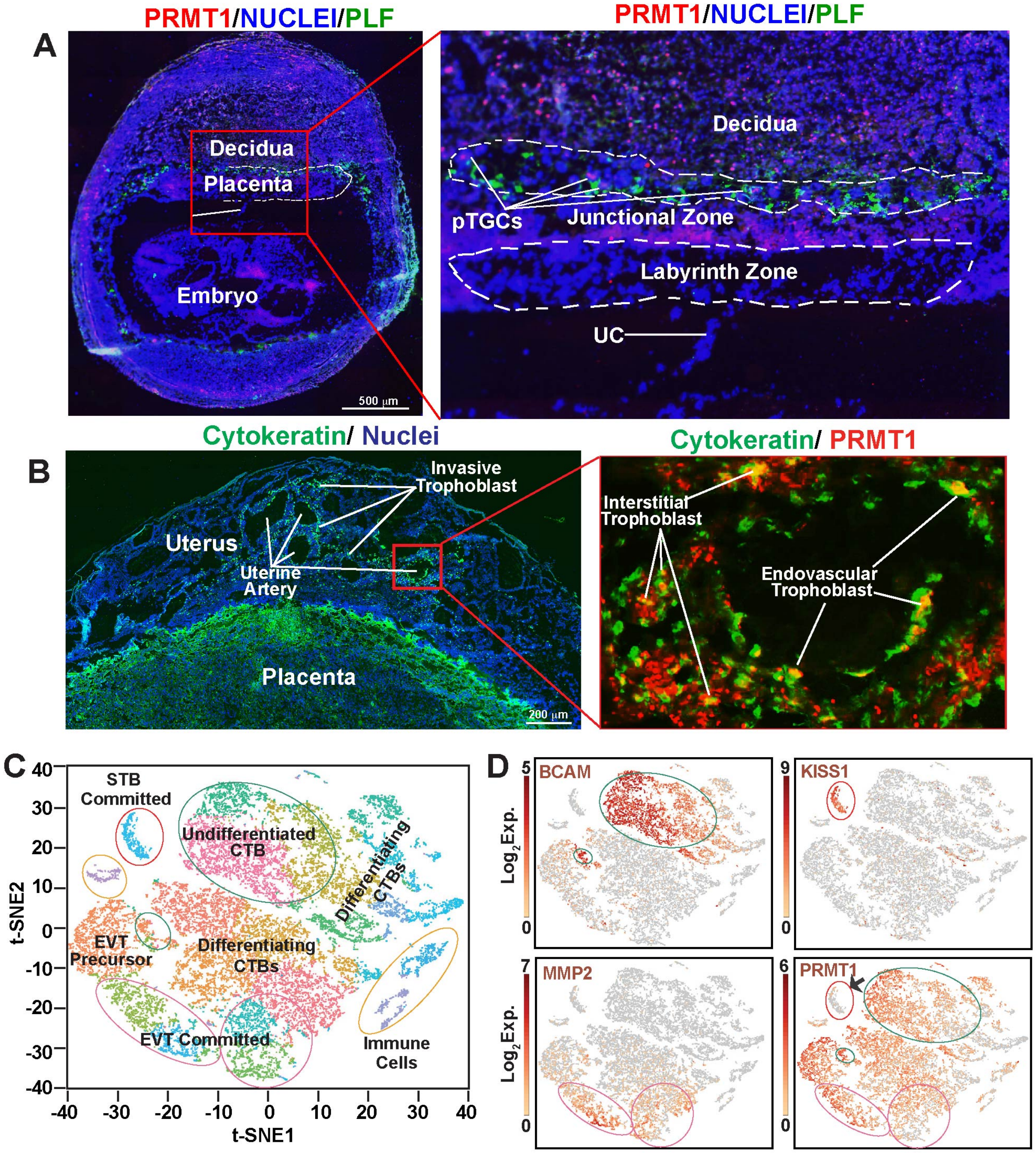


**Figure S2. (A)** Immunofluorescence analysis of PRMT1 protein expression in a histological section of mouse E9.5 conceptus. The inset shows that PRMT1 protein is mostly undetectable in the developing labyrinth but is detectable in the junctional zone. **(B)** Immunostaining of an E18.5 rat uterine-placental interface showing PRMT1 expression in both interstitial and endovascular invasive trophoblast cells. **(C)** scRNA-seq analyses in first-trimester human placentae. The t-SNE plot of cell clusters showing distinct trophoblast and non-trophoblast cells. **(D)** Expressions of specific genes are shown to identify different trophoblast clusters. BCAM for undifferentiated CTBs, KISS1 for CTBs that are committed for STB development. MMP2 for commited EVT precursors. Note that PRMT1 expression is repressed in committed STBs.

**Fig. S3**

**A**

| Analyses in Global *Prmt1* knockout (*Prmt1* KO) mouse embryos | | |
|---|---|---|
| **Embryo Genotype** | **Number of Embryos** | **Embryos with Defective Placenta Primordium** |
| **Wild Type** | **14** | **0** |
| ***Prmt1*** $^{+/-}$ | **25** | **0** |
| ***Prmt1* KO** | **12** | **12** |

**B**

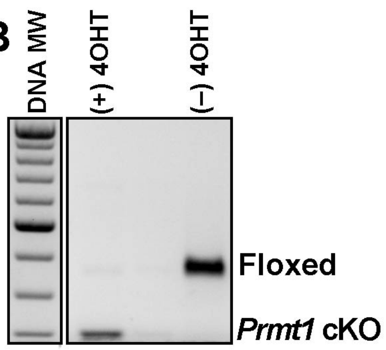


**C**

| Analyses in *Prmt1* conditional knockout (*Prmt1* cKO) mouse embryos | | | | |
|---|---|---|---|---|
| **Dev. Stage** | **Conceptus analyzed (Total embryos 52)** | | **Embryos with developmental defects** | |
| **E7.5** | **Floxed Control** | ***Prmt1* cKO** | **Floxed Control** | ***Prmt1* cKO** |
| | **36** | **16** | **0** | **16** |

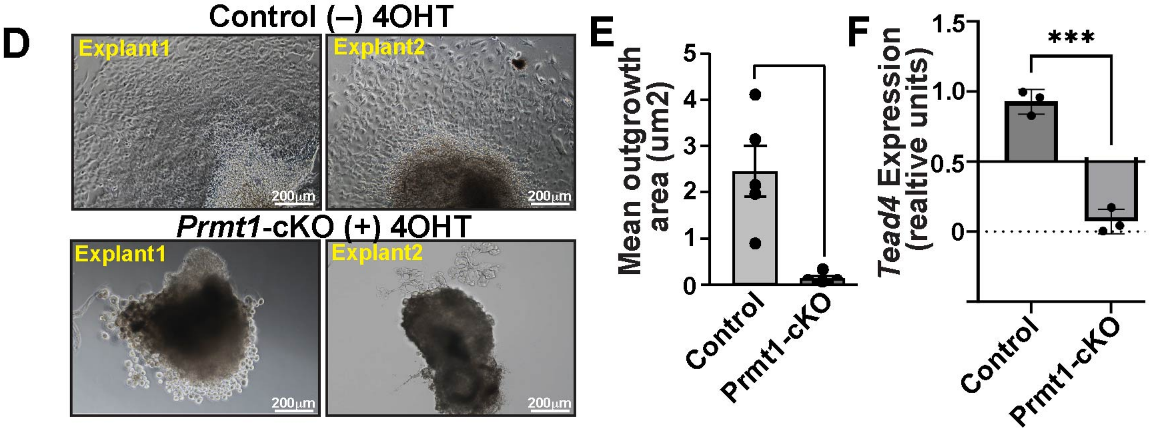


**Figure S3**. **(A)** Table showing number and phenotype of Wild-type, *Prmt1* $^{(+)/(-)}$ and *Prmt1-KO* embryos, analyzed ~ E7.0. 5 independent experiments with multiple pregnant females were conducted. **(B)** Genotype analyses confirming *Prmt1* $^{fl/fl}$ and *Prmt1-cKO* mouse embryos. DNA MW, molecular weight marker. **(C)** Table showing number and phenotype of *Prmt1* $^{fl/fl}$ and *Prmt1-cKO* embryos, analyzed ~ E7.5. 6 independent experiments were conducted. **(D)** E7.5 mouse placenta primordia containing the ExE/EPC regions were isolated for testing the importance of PRMT1 in primary TSPC self-renewal. *Prmt1* $^{fl/fl}$ (Control) and *Prmt1-cKO* ExE/EPC explants were cultured with Tamoxifen in FGF4/Heparin containing mouse TSC culture medium. Micrographs show severe defect in expansion of primary TSPCs upon *Prmt1* deletion. **(E)** Quantitative analyses of outgrowths of explant cultures of ExE/EPC regions from control and *Prmt1-cKO* embryos (mean ± SE; n=5 ExE/EPC cultures for each condition, $p \leq 0.05$). **(F)** RT-qPCR in control and *Prmt1-cKO* ExE/EPC showing loss of *Tead4* transcription in TSPCs upon *Prmt1* deletion.

Fig. S4

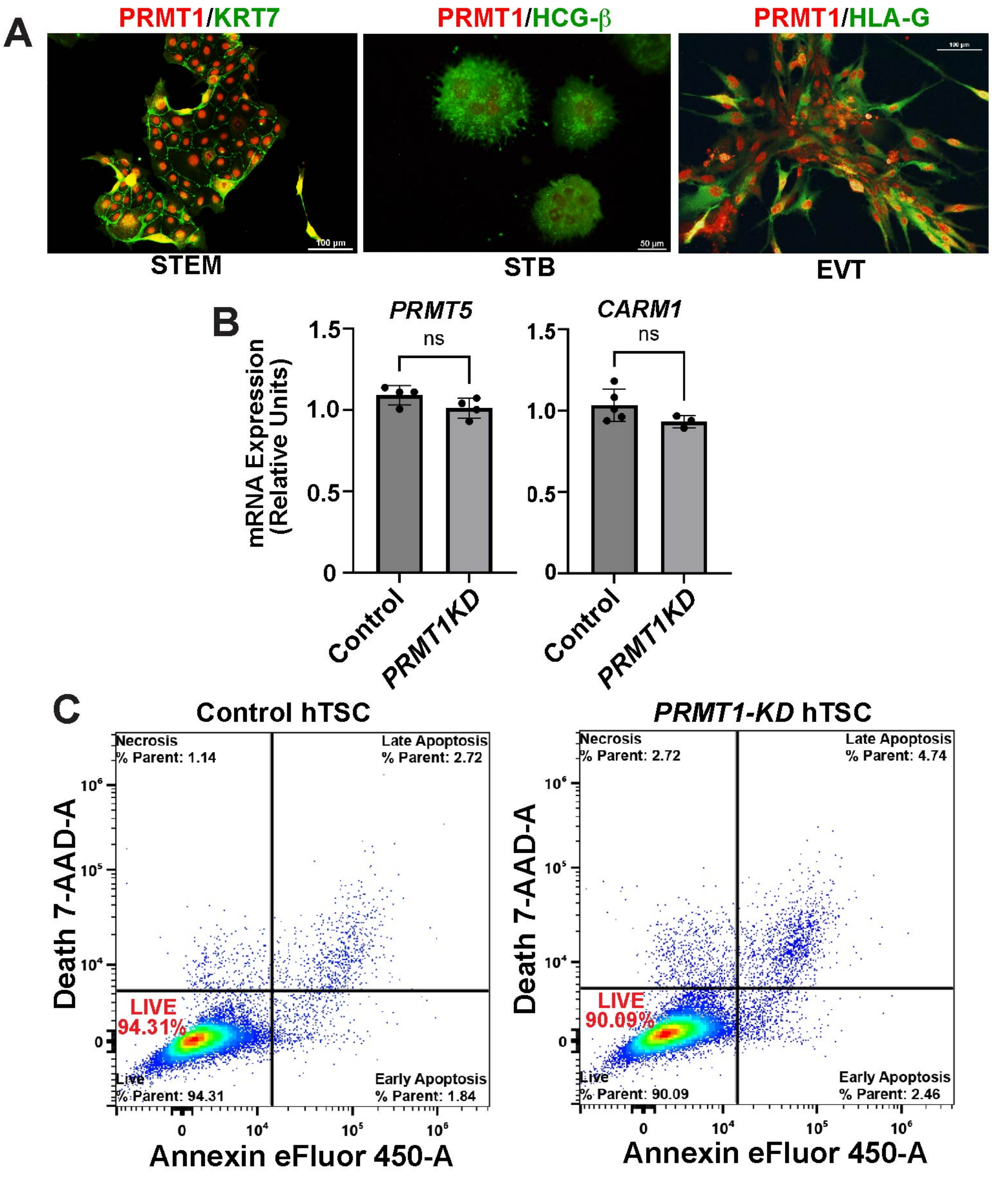


**Figure S4. (A)** Immunofluroescence images showing PRMT1 protein expression in undifferentiated hTSCs and after their differentiation to STBs and EVTs. Note loss of PRMT1 expression in STBs. **(B)** RT-qPCR analyses of PRMT5 and CARM1 mRNA expression in control and PRMT1-KD hTSCs (n= 4 independent experiments). **(C)** Flow-cytometry analyses showing that depletion of PRMT1 does not induce cell death in hTSCs.

Fig. S5

A

| Repressed TSC Genes | Activated STB Genes | Repressed EVT Genes |
|---|---|---|
| *ATF2* | *CGA* | *NOTUM* |
| *MPI* | *CGB* | *MMP11* |
| *MYBL2* | *ERVW1* | *PLAC8* |
| *NUBPL* | *ESRRG* | *DLC1* |
| *SMAD5* | *FOXI3* | *IFI27* |
| *TEAD4* | *GCM1* | *AHDC1* |
| *TP63* | *OVOL1* | *EXTL3* |
| | *PRKCZ* | *EFNB1* |
| | *PSG4* | *EDNRB* |
| | *TFEB* | *PIPOX* |

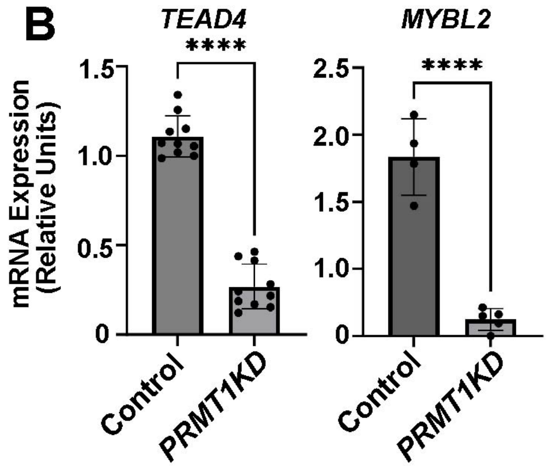


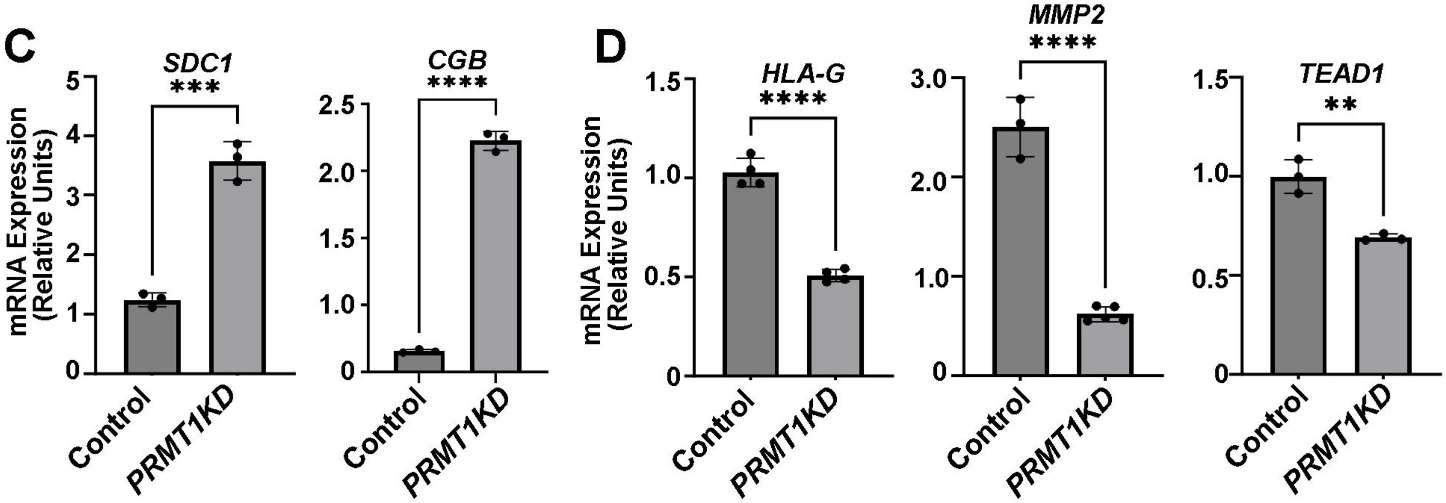


**Figure S5**. **(A)** Global RNA-seq analyses in *PRMT1-KD* hTSCs identified that TSC stem state and EVT-specific genes were downregulated and STB-specific genes were upregulated. The table shows examples of key genes of each cell type. **(B)** RT-qPCR confirming downregulation of *TEAD4* and *MYBL2* mRNA expression in *PRMT1-KD* hTSCs (n=at least 4 independent experiments were performed for each gene, p≤0.01). **(C)** RT-qPCR confirming upregulation of STB-specific genes *SDC1* and *CGB* in *PRMT1-KD* hTSCs, when cultured in stem state culture condition (n= 3independent experiments were performed, p≤0.01). (D) RT-qPCR showing inefficient induction of EVT-specific genes in *PRMT1-KD* hTSCs, when cultured in EVT differentiation condition (n= at least 3 independent experiments were performed for each gene, p≤0.01).

**Fig. S6**

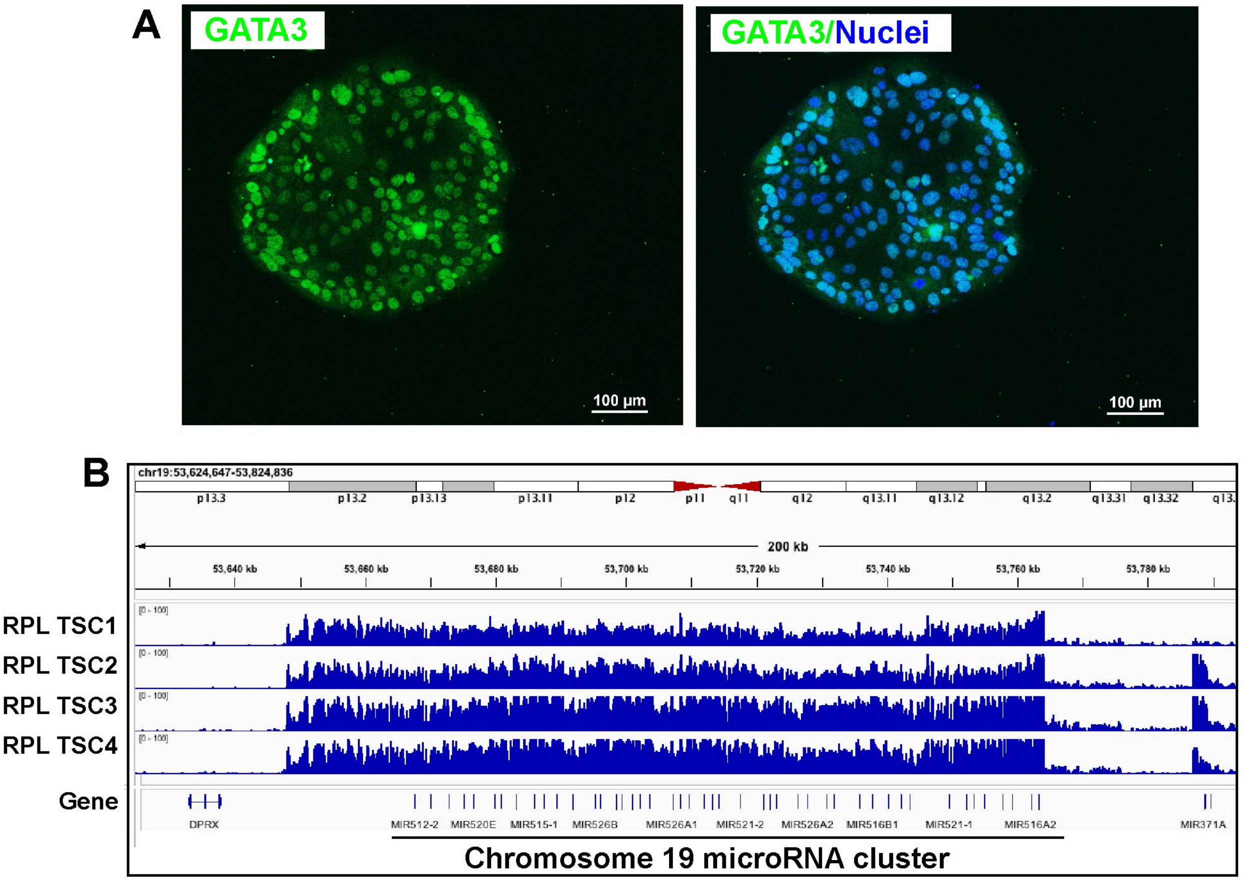


**Fig. S6 (A)** Immunofluorescence Images confirming GATA3 Expression in RPL-TSCs. **(B)** RNA-seq analyses confirming high expressions of chromosome 19 micro RNAs in RPL-TSCs.

**Fig. S7**

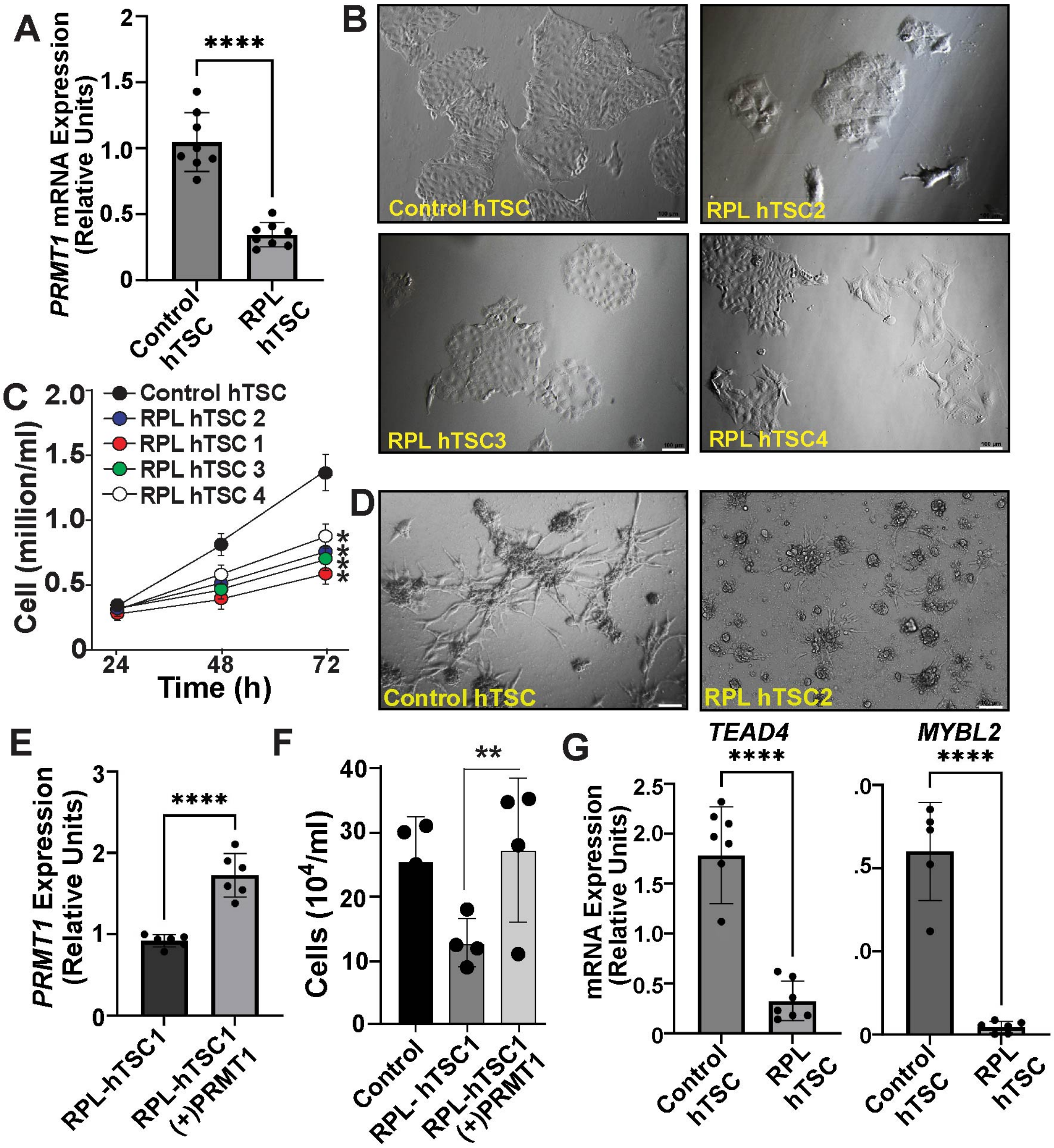


**Figure S7. (A)** RT-qPCR showing loss of *PRMT1* mRNA expression in RPL-hTSCs. Two control hTSC lines, CT27 and CT29, were used and four RPL-hTSC lines were analyzed. p≤0.001. **(B)** Representative colony morphology of control CT27 hTSCs and four RPL-hTSCs after 72h in culture in stem state culture condition. Note smaller colony sizes of RPL-hTSC lines. **(C)** Equal numbers of control CT27 and RPL-hTSCs were cultured and cell numbers were counted over 72h after seeding. The plot shows reduced cell proliferaion rate of RPL-hTSC lines. (n= Three independent experiments, p≤0.01). **(D)** Representative micrograps showing defective EVT differentiation of an RPL-hTSC line when cultured in EVT differentiation condition for 6 days. CT27 hTSCs were used as control. **(E)** RT-qPCR showing rescue of *PRMT1* mRNA expression in a RPL-hTSC line. (n= Five independent experiments, p≤0.01). **(F)** The plot shows increase in cell proliferaion rate of an RPL-hTSC line after rescue of PRMT1 expression. (n= Three independent experiments, p≤0.01). **(G)** RT-qPCR showing rescue of *TEAD4 and MYBL2* mRNA expression in a RPL-hTSC line after rescue of PRMT1 expression. (n= at least six independent experiments for each gene, p≤0.001).

### Figure S8

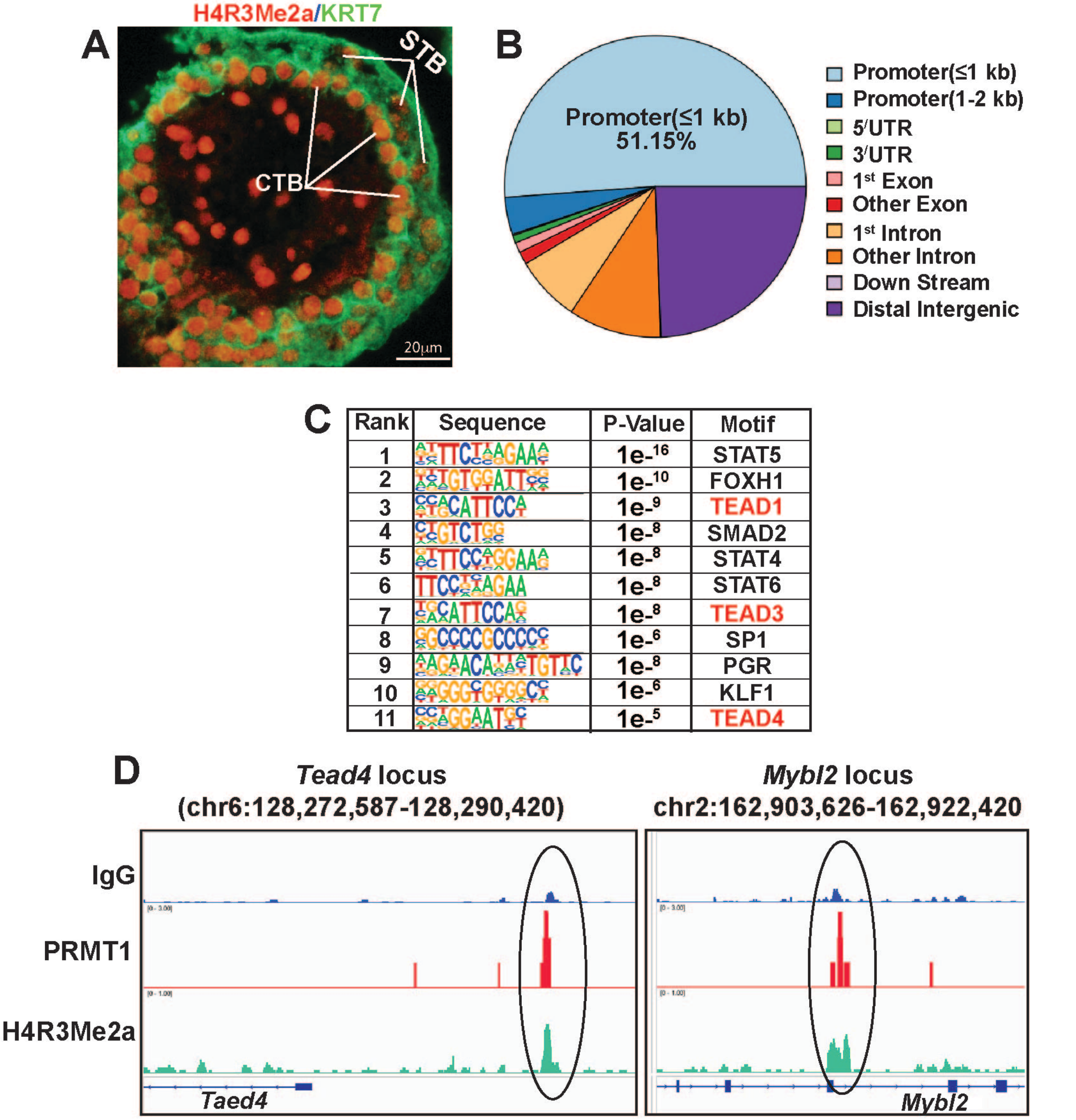


**Figure S8. (A)** Immunofluorescence image of a cross-section of a first-trimester (7+3 week) human placenta showing relatively high H4R3Me2a modification in CTBs compared to STBs. (B) The Pie chart shows the distribution of PRMT1 chromatin binding regions in hTSCs. More than 50% PRMT1 chromatin binding regions are within 1 kb of gene promoters. (C) HOMER motif analyses showing most significantly enriched transcription factor binding motifs at the PRMT1 chromatin binding regions. Note prevalance of TEAD and STAT motifs in PRMT1 binding regions. (D) IGV tracks showing PRMT1 binding peaks and H4R3Me2a enrichment at the *Tead4* and *Mybl2* loci in TSPCs of E7.5 ExE/EPC region.

#### Figure S9

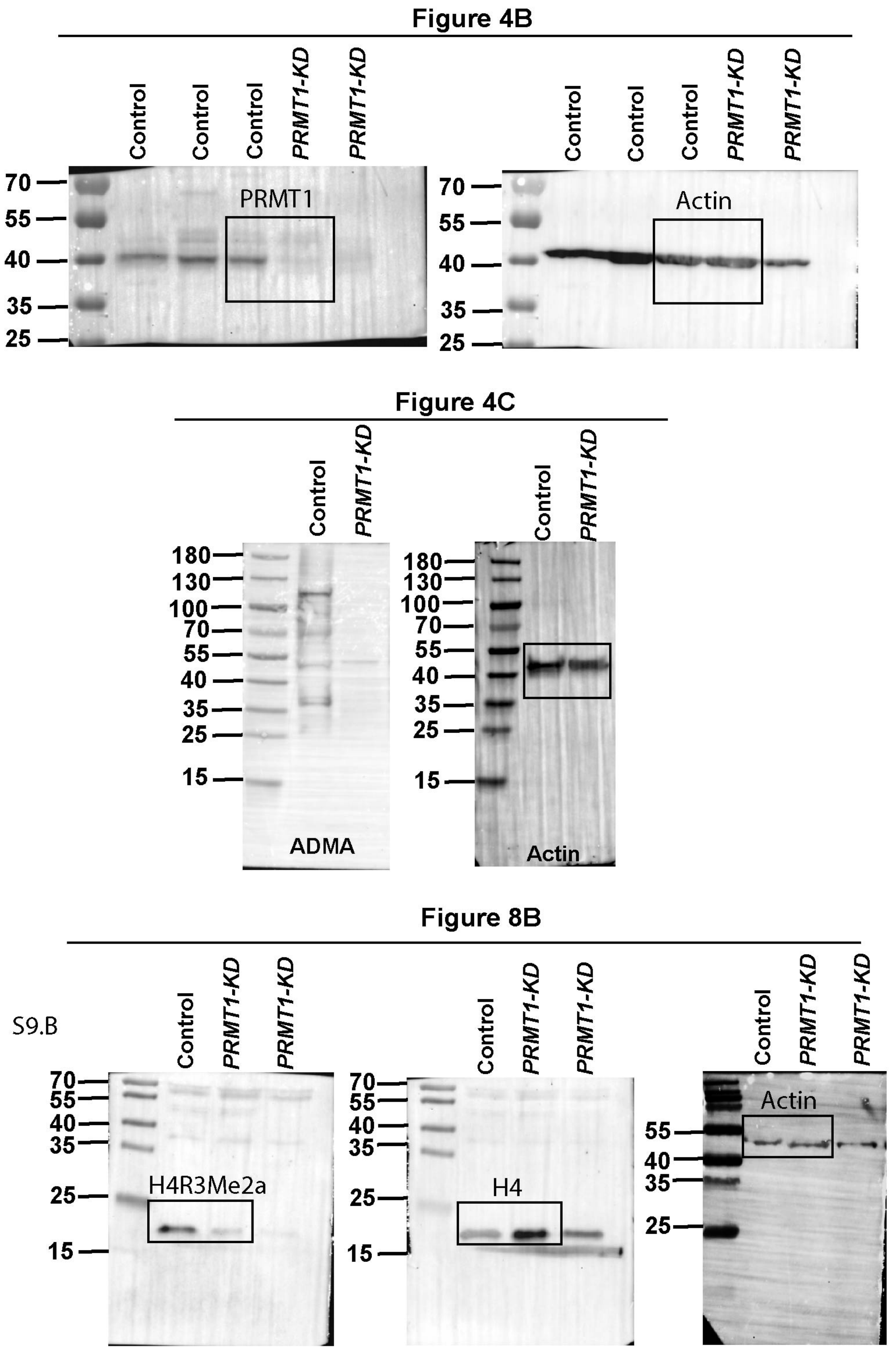


**Figure S9: Original Western blots that are used for different figures.**

**Supplementary Table 1: Idiopathic RPL Placenta Tissue Used For The Study**

| Sample No | Gestational age (Days) | Placental Villi Defect | Age of Mother | ≥ 2 prior miscarriages |
|---|---|---|---|---|
| 1 | 42 | Yes | 31 | Yes |
| 3 | 44 | No | 29 | Yes |
| 3 | 42 | No | 27 | Yes |
| 4 | 104 | Yes | 33 | Yes |
| 5 | 45 | No | 26 | Yes |
| 6 | 47 | No | 32 | Yes |
| 7 | 43 | No | 30 | Yes |
| 8 | 46 | Yes | 28 | Yes |
| 9 | 87 | Yes | 32 | Yes |
| 10 | 56 | No | 33 | Yes |
| 11 | 63 | Yes | 36 | Yes |
| 12 | 56 | Yes | 27 | Yes |

**Supplementary Table 2: Genotyping Primers**

| | | |
|---|---|---|
| *Prmt1-Tm1a* | TGGATGGAGGATGGACAGTG | GAACTTCGGAATAGGAACTTCG |
| *Prmt1-Flox* | TGGATGGAGGATGGACAGTG | CGAGTAGCAAGGAGGTCGAT |
| PRMT1-KO | AAGGCGCATAACGATACCAC | ACTGATGGCGAGCTCAGACC |
| *Cre* | AAAATTTGCCTGCATTACCG | ATTCTCCCACCGTCAGTACG |
| *Flp* | GCGAAGAGTTTGTCCTCAACC | AAAGTCGCTCTGAGTTGTTAT |

**Supplementary Table 3: Antibodies Used**

| Primary Antibody | Company | Catalog Number |
|---|---|---|
| PRMT1 | Cell SignalingTechnology | 2449 |
| H4R3me2a | Active Motif | 39705 |
| Pan-cytokeratin | Abcam | 9377 |
| Cytokeratin 7 | Dako | M7018 |
| Esrrb | Thermo Fisher Scientific | PPH670500 |
| TEAD4 | Abcam | 58310 |
| β-Actin | Sigma | A5441 |
| MYBL2 | Proteintech | 18896-I-AP |
| H4 | Cell Signaling Technology | 2592S |
| HCGβ | Abcam | 53087 |
| HLA-G | Abcam | 7758 |
| E-cadherin | Abcam | 1416 |
| Ki67 | Abcam | 16667 |
| ADMA | Dr Bedford's Lab | |
| **Secondary Antibody (IgG)** | | |
| Alexa fluor 488 donkey antimouse IgG | Invitrogen | A21202 |
| Alexa fluor 568 donkey antirabbit IgG | Invitrogen | A10042 |
| Goat anti-mouse IgG-HRP | Santa Cruz | sc2005 |
| Goat anti-rabbit IgG-HRP | Santa Cruz | sc2004 |

**Supplementary Table 4: Primers Used for RT-qPCR**

| Name | Forward | Reverse |
|---|---|---|
| *18sRNA* | AACCCGTTGAACCCCATT | CCATCCAATCGGTAGTAGCG |
| *TEAD4* | ACG GCC TTC CAC AGT AGC AT | CTT GCC AAA ACC CTG AGA CT |
| *TP63* | GTCATTTGATTCGAGTAGAGG GG | CTGGGGTGGCTCATAAGGT |
| *PRMT1* | CTTTGACTCCTACGCACACTT | GTGCCGGTTATGAAACATGGA |
| *CARM1* | TGGGCTACATGCTCTTCAACG | CGTCACCAATGGTAGGAAACAT |
| *PRMT5* | CGATCAGACCTACTGCTGTCA | CTCGGAGTTCCTGCGAATCT |
| *SDC1* | CTATTCCCACGTCTCCAG AACC | CTATTCCCACGTCTCCAGAACC |
| *CGB* | GTGTGCATCACCGTCAACAC | GGTAGTTGCACACCACCTGA |
| *TP63* | GTCATTTGATTCGAGTAGAGGGG | CTGGGGTGGCTCATAAGGT |
| *MYBL2* | CTTGAGCGAGTCCAAAGACTG | AGTTGGTCAGAAGACTTCCCT |
| *H-LAG* | CCACCACCCTGTCTTTGACTAT | AGCTCCTGGGTCTGGTCCT |
| *TEAD1* | ATGGAAAGGATGAGTGACTCTGC | TCCCACATGGTGGATAGATAGC |
| *MMP2* | TCTCCTGACATTGACCTTGGC | CAGGGTGCTGGCTGAGTAGATC |
